# Brain-Age Prediction: Systematic Evaluation of Site Effects, and Sample Age Range and Size

**DOI:** 10.1101/2023.11.06.565917

**Authors:** Yuetong Yu, Hao-Qi Cui, Shalaila S. Haas, Faye New, Nicole Sanford, Kevin Yu, Denghuang Zhan, Guoyuan Yang, Jia-Hong Gao, Dongtao Wei, Jiang Qiu, Boris Bernhardt, Paul Thompson, Sophia Frangou, Ruiyang Ge, ENIGMA World Aging Center

**Author notes:** Corresponding author: Sophia Frangou, Icahn School of Medicine at Mount Sinai, 1425 Madison Avenue, New York, New York, 10029, USA. Equal Contribution (Yuetong Yu, Hao-Qi Cui). ENIGMA World Aging Center Authors Group.

## Abstract

Structural neuroimaging data have been used to compute an estimate of the biological age of the brain (brain-age) which has been associated with other biologically and behaviorally meaningful measures of brain development and aging. The ongoing research interest in brain-age has highlighted the need for robust and publicly available brain-age models pre-trained on data from large samples of healthy individuals. To address this need we have previously released a developmental brain-age model. Here we expand this work to develop, empirically validate, and disseminate a pre-trained brain-age model to cover most of the human lifespan. To achieve this, we selected the best-performing model after systematically examining the impact of site harmonization, age range, and sample size on brain-age prediction in a discovery sample of brain morphometric measures from 35,683 healthy individuals (age range: 5-90 years; 53.59% female). The pre-trained models were tested for cross-dataset generalizability in an independent sample comprising 2,101 healthy individuals (age range: 8-80 years; 55.35% female) and for longitudinal consistency in a further sample comprising 377 healthy individuals (age range: 9-25 years; 49.87% female). This empirical examination yielded the following findings: (1) the accuracy of age prediction from morphometry data was higher when no site harmonization was applied; (2) dividing the discovery sample into two age-bins (5-40 years and 40-90 years) provided a better balance between model accuracy and explained age variance than other alternatives; (3) model accuracy for brain-age prediction plateaued at a sample size exceeding 1,600 participants. These findings have been incorporated into CentileBrain [https://centilebrain.org/#/brainAGE2], an open-science, web-based platform for individualized neuroimaging metrics.

## 1. INTRODUCTION

Prior literature has documented extensive age-related changes in brain morphology as inferred from structural magnetic resonance imaging (sMRI) studies [Hogstrom et al., 2013; Bethlehem et al., 2022; Dima et al., 2022; Frangou et al., 2022; Ge et al., 2023]. Machine learning algorithms can model these age-related changes to generate an estimate of the biological age of the brain (brain-age) [Baecker et al., 2021; Schulz et al., 2022; More et al., 2023]. Brain-age estimates derived from healthy individuals can be used to establish a normative reference pattern for typical development and aging. In each individual, large deviations between brain-age and chronological age indicate atypical development or aging [Cole & Franke, 2017; Franke & Gaser, 2019; Ball et al., 2021; Modabbernia et al., 2022].

Key parameters that influence accuracy in any brain-age prediction workflow comprise the type of morphometric input features and machine learning algorithms, the size and age range of the sample, and the handling of site-effects, in the case of pooled samples. Input features include voxel-wise data [Cole et al., 2020; Baeker et al., 2021], or data derived via dimensionality reduction through atlas-based parcellation [Modabbernia et al.,2022] or statistical methods (e.g., principal component analysis) [Franke et al., 2013]. Generally, there is no advantage to using voxel-wise data or highly granulated parcels [Valizadeh et al., 2017; Baeker et al., 2021; Modabbernia et al., 2022]. There are also multiple algorithms for computing brain-age that comprise conventional methods, such as linear and Bayesian models, tree-based and kernel-embedded models, and artificial neural networks commonly referred to as deep learning networks [Goodfellow et al., 2016]. Studies that have undertaken a comparative evaluation of these algorithms on the accuracy of sMRI-derived brain-age estimates collectively suggest that conventional methods outperform deep learning networks in addition to being computationally more efficient [Couvy-Duschesne et al., 2020; He et al., 2020; de Lange et al., 2022; Grinstain et al., 2022; Modabbernia et al., 2022; More et al., 2023].

We have previously shown that Support Vector Regression (SVR) with Radial Basis Function (RBF) Kernel is preferable to parametric and nonparametric, Bayesian, linear and nonlinear, and other kernel-based models particularly because of its resilience to extreme outliers [Modabbernia et al. 2022]. We adopted this algorithm to build a developmental brain-age model based on morphometric data from healthy youth aged 5-22 years [Modabbernia et al., 2022] and made this freely available to the scientific community through a web platform dedicated to providing models for individual-level neuroimaging measures [https://centilebrain.org/#/brainAGE]. Here we extend our previous work to construct brain-age prediction models that are empirically validated and provide greater coverage of the human lifespan. To achieve this, we pooled brain morphometric data from 35,683 healthy individuals (aged 5-90 years), as the discovery sample, and data from an independent sample totaling 2,102 healthy individuals (aged 27.74 years, as the replication sample. We evaluated the effect of age and sample composition on model performance as there is no consensus regarding the optimal method for integrating these parameters into brain-age models. It is acknowledged that site harmonization strategies [Lombardi et al., 2020] significantly affect the performance of brain-age models. Moreover, brain-age studies have focused either on youth [Ball et al., 2021; Brower et al., 2021; Luna et al., 2021] or on middle-aged and elderly individuals [Cole & Franke, 2017; Elliot et al., 2021]. Thus, the workflow required for reliable brain-age estimates in samples that cover most of the lifespan remains unclear. To address these knowledge gaps, we empirically evaluated the performance of the SVR-RBF algorithm in our discovery sample using diverse site harmonization strategies and by resampling the discovery model to produce subsets of different sizes and age ranges. The resulting models were then tested on the replication sample for cross-sample performance and longitudinal consistency. We outline our method in detail while codes and the best-performing models are freely available on our dedicated web platform [https://centilebrain.org/#/brainAGE2].

## 2. METHODS

### 2.1. Samples

Different independent samples were used for discovery, replication, and longitudinal consistency. These samples included pooled multisite sMRI data from Australia, East Asia, Europe, and North America (supplementary section S1, Figure S1). The discovery sample comprised 35,683 healthy individuals (53.59% female, age range 5-90 years; Table S1). The replication sample comprised a total of 2101 healthy individuals (55.35% female, age range 8-80 years; Table S2). The longitudinal consistency sample included data from 377 healthy individuals (age range: 9-25 years; 49.87% female; Table S2) participating in the Southwest Longitudinal Imaging Multimodal Study (SLIM) and the Queensland Twin Adolescent Brain Study (QTAB). Only high-quality morphometric measures (supplementary section S2) were included from participants who were free of psychiatric, medical, and neurological morbidity and cognitive impairment at the time of scanning.

### 2.2. Brain Morphometric Input Features

Morphometric feature extraction from whole-brain T1-weighted images was implemented using the standard pipelines in the FreeSurfer image analysis suite (http://surfer.nmr.mgh.harvard.edu/) to yield a total of 150 morphometric features that have been extensively utilized in prior models for predicting brain age [Elliott et al., 2021; Han et al.2021; de Lange et al.,2022]. These comprised Desikan-Killiany atlas measures of cortical thickness (n=68), cortical surface area (n=68) [Desikan et al., 2006], and regional subcortical volumes (n=14) based on the Aseg atlas [Fischl et al., 2002].

### 2.3. Evaluation of Brain-Age Models

#### 2.3.1. Core Elements

i. All brain-age models evaluated were sex-specific because of the known sex differences in brain morphometry. The method of evaluation was identical for both sexes.
ii. All models used the same 150 input features described above.
iii. All models used the SVR-RBF which we adopted as our algorithm of choice as we have demonstrated its favorable performance in terms of accuracy, computational efficiency, and robustness to outliers when compared to other machine learning algorithms [Modabbernia et al., 2022]. This choice is supported by independent studies that have undertaken a comparative evaluation of multiple algorithms [Beheshti et al., 2022; More et al., 2023].
iv. The primary performance measures for all models were the mean absolute error (MAE), which represents the absolute difference between brain-age and chronological age, and the correlation coefficient (CORR) between brain-age and chronological age.
v. Brain-age is often overestimated in younger individuals and underestimated in older people [Liang et al., 2019; de Lange & Cole, 2020]. To counter this bias, we implemented a robust approach to adjust this age-related bias following Beheshti and colleagues [Beheshti et al., 2019]. However, as age bias-corrected metrics often reflect elevated accuracy, even for models with poor performance [de Lange et al., 2022], we focus primarily on uncorrected model performance.

#### 2.3.2. Analysis Workflow

The procedures used to generate optimized sex-specific models are illustrated in Figure 1. For all models, hyper-parameter tuning (C and sigma) was performed in the discovery sample using a grid search approach in a 10-fold cross-validation scheme across five repetitions. In each cross-validation, 90% of the discovery sample was used to train the model and 10% was used to test the model parameters; subsequently, the model was retrained on the entire discovery sample using the optimal hyperparameters identified from the cross-validation process. As detailed in the subsequent sections, first, we tested three different strategies for handling site effects. The site-harmonization strategy used in the subsequent procedures was selected based on its superior performance, as indicated by the lowest within-sample cross-validation MAE (MAE_CV_) and the highest CORR (CORR_CV_) in the discovery sample. The model with the lowest replication MAE and highest replication CORR in the replication sample (referred to MAE_R_ and CORR_R_) and in the longitudinal consistency samples was chosen as the preferred model.

**Figure 1.**
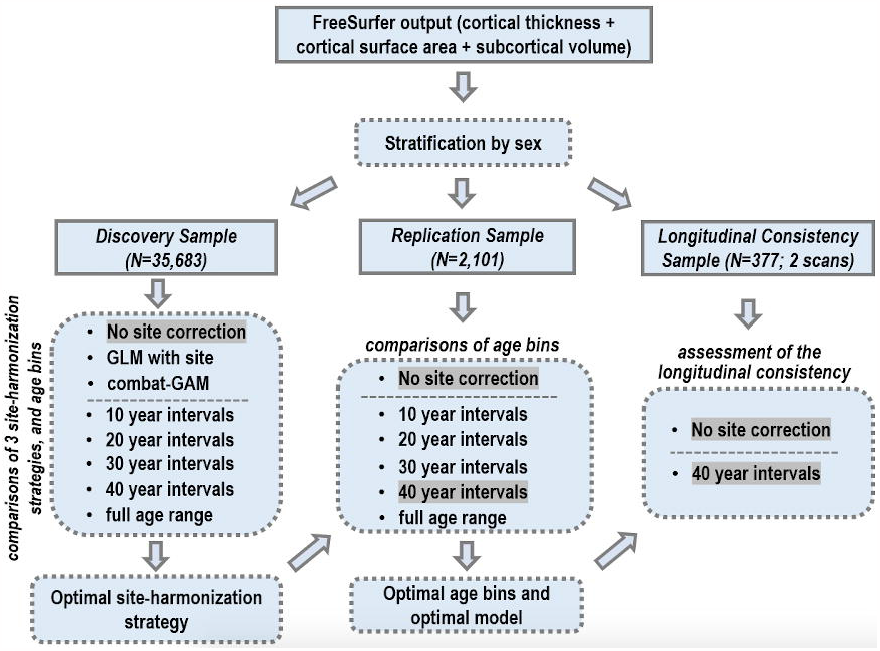
Flowchart of brain age model optimization: after conducting the analysis with FreeSurfer and stratifying the samples by sex, the study proceeded as follows. (1) The discovery sample was utilized to evaluate the impact of site-harmonization strategies and age range. This analysis yielded the optimal site-harmonization strategy, and this optimal site-harmonization strategy was highlighted with a gray background. (2) The independent replication sample was employed to further investigate the influence of age range. The outcome of this analysis led to the determination of the optimal age bins and final models, with the chosen optimal age bins being highlighted in gray. (3) The independent longitudinal consistency sample was utilized to assess the longitudinal consistency of the pre-trained optimal models.

#### 2.3.3. Evaluation of Site Effects and Age Range in the Discovery Sample

We evaluated three site handling strategies in each of five scenarios after partitioning the discovery sample into different age bins as follows: (i) a single bin with the full sample age range (5-90 years); (ii) nine bins each covering sequential 10-year intervals, i.e., age<10 years, 10≤age<20 years, 20≤age<30 years, 30≤age<40 years, 40≤age<50 years, 50≤age<60 years, 60≤age<70 years, 70≤age<80 years, and 80≤age<90 years; (iii) four bins each covering sequential 20-year intervals, i.e., age<20 years, 20≤age<40 years, 40≤age<60 years, and 60≤age≤80 years; (iv) three bins each covering sequential 30-year intervals i.e., age<30 years, 30≤age<60 years, and 60≤age≤90 years; (v) two age bins each covering sequential 40-year intervals, i.e., age<40 years, and 40≤age≤80 years. The following three site handling strategies were applied to each bin: (i) data residualisation with respect to the scanning site using Combat-GAM [Pomponio et al., 2020]; (ii) data residualization with respect to the scanning site using a generalized linear model [de Lange et al., 2022], and (iii) no site harmonization. The approach and age partition with the best-performing MAE_CV_ and CORR_CV_ values were considered for further evaluation.

#### 2.3.4. Evaluation of Site Effects and Age Range in the Replication Sample

The replication sample was partitioned in age bins similarly to the discovery sample and the pre-trained models were applied. The age bin partition that yielded the highest performing MAE_R_ and CORR_R_ values was identified as the preferred age bins.

#### 2.3.5. Estimation of the Minimum Sample Size

Subsequently, the discovery sample was randomly partitioned into 30 sex-specific subsets, ranging from 200 to 6,000 participants in increments of 200, without replacement. The robustness of the optimised sex-specific models to sample size in terms of CORR_CV_ and MAE_CV_ was assessed in each partition using 10-fold cross-validation with five repetitions. This analysis was performed individually for each of the preferred age bins according to section 2.3.4.

#### 2.3.6. Longitudinal Consistency

The longitudinal consistency sample included T1-weighted scans from a total of 377 participants scanned twice with an average interval of 1.89 (0.56) years. This sample was also divided into the preferred age bins as in the discovery and replication samples in section 2.3.4. The percentage change in MAE and CORR between the second scan and the first scan was evaluated in each age bin.

## 3. RESULTS

### 3.1. Site and Sample Age Range

Figure 2 illustrates cross-validated model performance within the distinct age bins of the discovery sample and Figure 3 illustrates the results of employing these pre-trained models in the independent replication sample. For simplicity, both figures display the results averaged across sexes as the sex-specific results were identical for males and females (Figures S2-S4, and Table S3-S6). For both sexes, omitting any site correction consistently demonstrated superior performance in terms of attaining the lowest MAE_CV_ values and highest CORR_CV_ values across the age bins.

**Figure 2.**
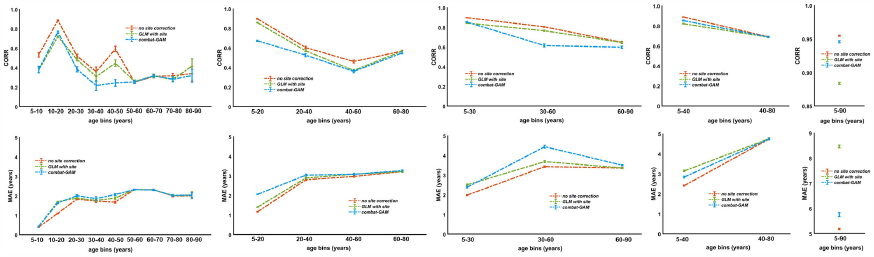
Performance metrics derived from repeated cross-validation in different age bins of the discovery sample. Each line represents one of the three site handling methods: Red=no site correction; Blue=site harmonization with Combat-GAM; Green=site data residualization using a generalized linear model (GLM); CORR, correlation coefficient between brain-age and chronological age; MAE: mean absolute error between brain-age and chronological age. Sex-specific results in Supplementary Figures S1 and S2.

**Figure 3.**
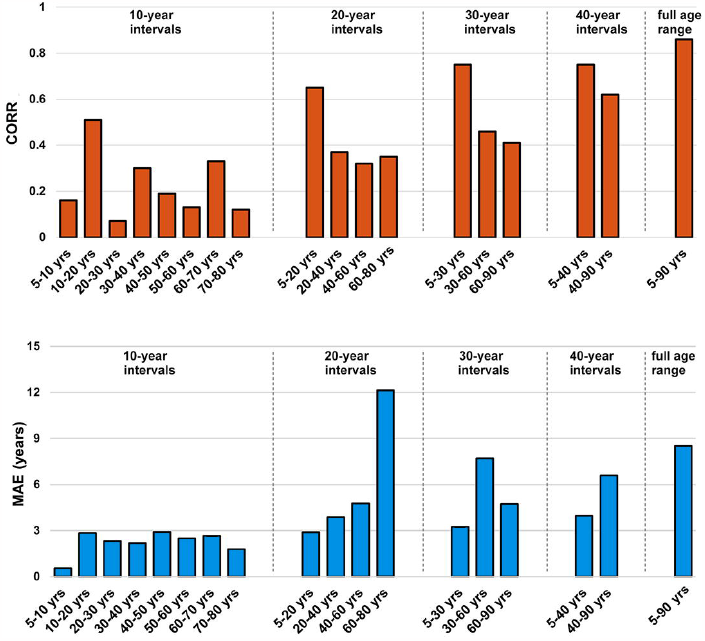
Performance metrics derived from the application of the models pre-trained on different age bins of the discovery sample to the corresponding age bins of the replication sample. CORR values averaged across sexes were 0.23 for 10-year interval bins; 0.42 for 20-year interval bins; 0.54 for 30-year interval bins; 0.68 for 40-year interval bins; and 0.86 for the full age range of the sample. MAE values averaged across sexes were 2.21 years for 10-year interval bins; 5.92 years for 20-year interval bins, 5.22 years for 30-year interval bins; 5.28 years for 40-year interval bins; and 8.52 years for the full age range of the sample. Sex-specific results are presented in supplementary Figure S4. CORR: correlation coefficient between brain-age and chronological age; MAE: mean absolute error between brain-age and chronological age.

Both in the discovery and the replication sample, the CORR and MAE were generally higher in age bins that included a wider age range. In other words, models based on a wider age range accounted for more of the variance in age but were less accurate. Therefore, in order to achieve a balance between CORR and MAE, we selected the two-bin partition with sequential 40-year intervals (i.e., 5-40 years and 40-90 years). By adopting this approach, we managed to attain a relatively low MAE while maintaining a relatively high CORR across sexes and age bins. Specifically, the respective average MAE_CV_ and CORR_CV_ were 3.55 (1.17) years and 0.79 (0.10) (Figures S2-S3, Tables S3-S4) and the average MAE_R_ and CORR_R_ were 5.28 years and 0.68 (Figure S4-S5, Tables S5-S6).

On average, age-bias adjustment improved the CORR_CV_ and MAE_CV_ by 79.67% and 35.56%, respectively in the discovery sample (Table S3 and Table S4) and improved the CORR_R_ and MAE_R_ by 287.06% and 41.79% in the replication sample (Table S5 and Table S6).

### 3.3. Effect of Sample Size

Figure 4 illustrates the effect of sample size in the discovery and replication samples using pre-trained models that were tested in 30 sex-specific subsets, ranging from 200 to 6,000 participants in increments of 200, without replacement. In the discovery sample, the CORR_CV_ improved in line with sample increase up to a size of 1,600 participants and it plateaued thereafter; the MAE_CV_ on the other hand exhibited smaller variation across varying sample sizes and stabilized around 1,000 participants (Figure S6 for sex-specific results). Similarly, in the replication sample, the CORR_R_ increased and MAE_R_ decreased as a function of the sample size until it reached 1,600 participants and plateaued thereafter.

**Figure 4.**
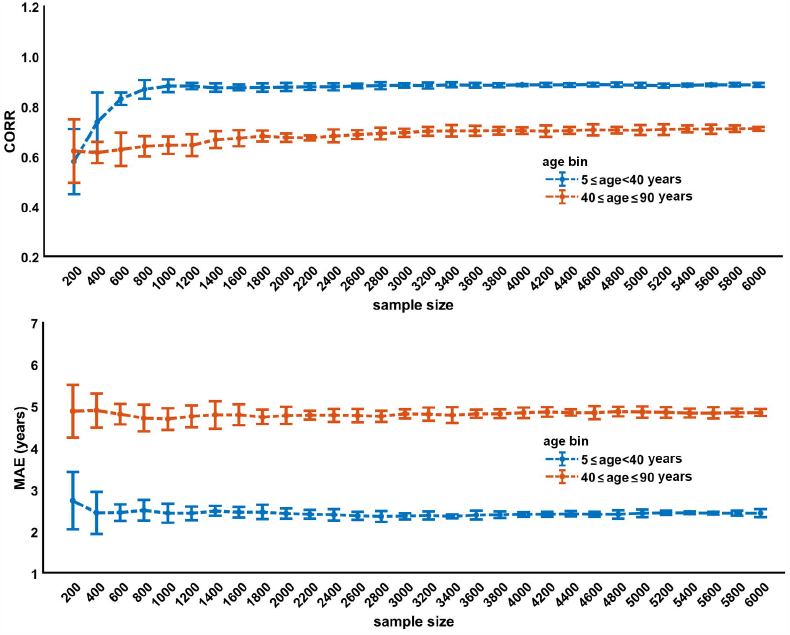
Model performance as a function of the discovery sample size in the two age bins (5-40 years and 40-90 years) of the discovery sample (left column) and replication sample (right column). Model parameters for each bin were obtained by randomly resampling the discovery sample without replacement generating subsets of 200-6,000 participants. The results are shown here as averages across sexes and the sex-specific findings are presented in supplementary Figure S6. CORR: correlation coefficient between brain-age and chronological age; MAE: mean absolute error between brain-age and chronological age.

### 3.4. Longitudinal Consistency

Figure 5 illustrates the stability of the pre-trained models in each age bin using the longitudinal consistency sample. The results indicated that models utilizing the two-bin partition (i.e., 5-40 years and 40-90 years) achieved optimal consistency on the longitudinal data. Sex-specific results are shown in supplementary Figures S7-S8 and Table S7-S8. On average, age-bias adjustment improved the CORR and MAE by 63.50% and 30.54%, respectively in the first scan of the longitudinal consistency sample; and age-bias adjustment improved the CORR and MAE by 73.39% and 20.87%, respectively in the second scan of the longitudinal consistency sample (Table S7 and Table S8).

**Figure 5.**
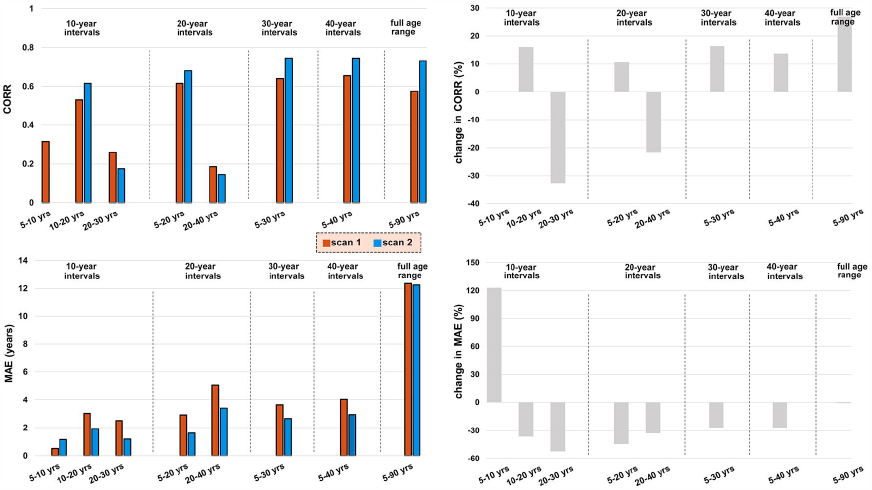
Model performance in longitudinal data. The left panel presents the CORR and MAE values for the first and second MRI scans, while the right panel exhibits the percentage changes (%) in CORR and MAE for the second scan compared to the first scan. The results were generated by employing models that had been trained on discovery samples from each age range division and then applied to the longitudinal consistency sample. CORR: correlation coefficient between brain-age and chronological age; MAE: mean absolute error between brain age and chronological age.

### 3.4. Data and Model Availability

Information about data availability is provided in Tables S1 and S2. Our dedicated web portal freely provides the optimal model parameters to be applied to any user-specified dataset in the context of open science. In addition to the pre-trained sex-specific models, the website provides tutorials and codes (https://centilebrain.org/#/tutorial4).

## 4. DISCUSSION

There is increased emphasis on the potential translational value of individualized neuroimaging measures such as brain-age that can be used to track deviation from typical brain development and brain aging [Cole & Franke, 2017; Franke & Gaser, 2019; Ball et al., 2021; Modabbernia et al., 2022]. The literature on morphometry-derived brain-age models from healthy individuals shows performance heterogeneity that is predicated on methodological differences in the specific features used, the algorithm employed, the handling of site-effects, the sample size and age distribution. The aim of the current investigation was to provide a resource to be used as a normative reference for brain-age by the scientific community. Having such a resource accomplishes at least four important objectives. First, it enables harmonization of the methods and models available for brain-age computation across studies. Second, it empowers researchers who do not have access to large normative datasets to generate reliable brain-age estimates in their own datasets. Third, it supports rigor and reproducibility in brain-age research. Fourth, together with our developmental brain-age model [Moddabenia et al., 2022], also available through our web platform, it provides models that cover most of the human lifespan (5-90 years) thus meeting the needs of researchers working in development or aging.

Following a systematic empirical evaluation, we selected SVR-RBF as the key algorithm [Modabbernia et al., 2022], and in this study, we determined the optimal site handling method for our model as well as the optimal age distribution for brain-age computation across most of the lifespan. This detailed evaluation was necessary as multiple prior studies have shown that site harmonization strategies as well as sample age distribution and size can influence model performance [de Lange et al., 2022]. As in previous reports, we found an inverse association between the age range of a sample, the MAE of the model, and the coefficient of correlation between brain-age and chronological age [de Lange et al., 2022]. MAE is generally lower in samples with a narrower age range which is attributable to the minimization of errors when the predicted brain-age approximates the mean chronological age of a sample. Concomitantly, the correlation between brain-age and chronological age becomes lower the narrower the age range of a sample [de Lange et al., 2022]. Previous reports have also shown that model accuracy for brain-age is generally better with larger sample sizes [de Lange et al., 2022]. Here we confirm this observation, but we also show that this relationship plateaus in samples with over 1,600 participants. This finding is particularly useful for evaluating the robustness of other existing models and for planning future studies.

The model proposed here suggests that an optimal balance between MAE and CORR is achieved when the lifespan sample is partitioned into two sequential age bins, 5-40 years and 40-90 years. The age-bias corrected MAE and CORR values for females in the 5-40 years age bin were 3.53 and 0.83 respectively, and in the 40-90 years age bin they were 4.45 and 0.86 respectively (also Table S5). In males, the age-bias corrected MAE and CORR values for females in the 5-40 years age bin were 3.60 and 0.84 respectively, and in the 40-90 years age-bin, they were 4.09 and 0.87 respectively (also Table S6). These values are well within the range reported in other studies that have evaluated different computational approaches to brain-age in healthy individuals. For example, More and colleagues [More et al., 2023] reported a range of MAE values between 4-8 years.

We appreciate that brain morphometric features are not the only type of neuroimaging measures that can be used to derive brain-age estimates. Other studies have used neuroimaging data from different modalities [Goyal et al., 2019; Beck et al., 2021; Lund et al., 2022; Zhou et al., 2023] or combinations of modalities [Cole, 2020; Niu et al., 2021; Rokiki et al., 2021]. Although it is important for the field to have a range of options for computing brain-age that can accommodate a variety of scientific questions, the wide availability and relative ease of acquiring and extracting brain morphometric data contribute to the popularity and preponderance of brain-age studies that use such data.

In conclusion, we present empirically validated models for brain-age that can accommodate studies using data across most of the lifespan. We have outlined the methodological choices that have led to these models as well as their performance within and across samples as well as longitudinally.

## Notes

### Competing Interest Statement

The authors have declared no competing interest.

## REFERENCES

Baecker L, Dafflon J, da Costa PF, Garcia-Dias R, Vieira S, Scarpazza C, Calhoun VD, Sato JR, Mechelli A, Pinaya WHL. Brain age prediction: A comparison between machine learning models using region- and voxel-based morphometric data. Hum Brain Mapp. 2021;42(8):2332–2346. doi: 10.1002/hbm.25368.

Ball G, Kelly CE, Beare R, Seal ML. Individual variation underlying brain age estimates in typical development. Neuroimage. 2021;235:118036. doi: 10.1016/j.neuroimage.2021.118036.

Beck D, de Lange AG, Maximov II, Richard G, Andreassen OA, Nordvik JE, Westlye LT. White matter microstructure across the adult lifespan: A mixed longitudinal and cross-sectional study using advanced diffusion models and brain-age prediction. Neuroimage. 2021;224:117441. doi: 10.1016/j.neuroimage.2020.117441.

Beheshti I, Nugent S, Potvin O, Duchesne S. Bias-adjustment in neuroimaging-based brain age frameworks: A robust scheme. Neuroimage Clin. 2019;24:102063. doi: 10.1016/j.nicl.2019.102063.

Beheshti I, Ganaie MA, Paliwal V, Rastogi A, Razzak I, Tanveer M. Predicting Brain Age Using Machine Learning Algorithms: A Comprehensive Evaluation. IEEE J Biomed Health Inform. 2022;26(4):1432–1440. doi: 10.1109/JBHI.2021.3083187.

Bethlehem RAI, Seidlitz J, White SR, Vogel JW, Anderson KM, Adamson C, Adler S, Alexopoulos GS, Anagnostou E, Areces-Gonzalez A, Astle DE, Auyeung B, Ayub M, Bae J, Ball G, Baron-Cohen S, Beare R, Bedford SA, Benegal V, Beyer F, Blangero J, Blesa Cábez M, Boardman JP, Borzage M, Bosch-Bayard JF, Bourke N, Calhoun VD, Chakravarty MM, Chen C, Chertavian C, Chetelat G, Chong YS, Cole JH, Corvin A, Costantino M, Courchesne E, Crivello F, Cropley VL, Crosbie J, Crossley N, Delarue M, Delorme R, Desrivieres S, Devenyi GA, Di Biase MA, Dolan R, Donald KA, Donohoe G, Dunlop K, Edwards AD, Elison JT, Ellis CT, Elman JA, Eyler L, Fair DA, Feczko E, Fletcher PC, Fonagy P, Franz CE, Galan-Garcia L, Gholipour A, Giedd J, Gilmore JH, Glahn DC, Goodyer IM, Grant PE, Groenewold NA, Gunning FM, Gur RE, Gur RC, Hammill CF, Hansson O, Hedden T, Heinz A, Henson RN, Heuer K, Hoare J, Holla B, Holmes AJ, Holt R, Huang H, Im K, Ipser J, Jack CR Jr, Jackowski AP, Jia T, Johnson KA, Jones PB, Jones DT, Kahn RS, Karlsson H, Karlsson L, Kawashima R, Kelley EA, Kern S, Kim KW, Kitzbichler MG, Kremen WS, Lalonde F, Landeau B, Lee S, Lerch J, Lewis JD, Li J, Liao W, Liston C, Lombardo MV, Lv J, Lynch C, Mallard TT, Marcelis M, Markello RD, Mathias SR, Mazoyer B, McGuire P, Meaney MJ, Mechelli A, Medic N, Misic B, Morgan SE, Mothersill D, Nigg J, Ong MQW, Ortinau C, Ossenkoppele R, Ouyang M, Palaniyappan L, Paly L, Pan PM, Pantelis C, Park MM, Paus T, Pausova Z, Paz-Linares D, Pichet Binette A, Pierce K, Qian X, Qiu J, Qiu A, Raznahan A, Rittman T, Rodrigue A, Rollins CK, Romero-Garcia R, Ronan L, Rosenberg MD, Rowitch DH, Salum GA, Satterthwaite TD, Schaare HL, Schachar RJ, Schultz AP, Schumann G, Schöll M, Sharp D, Shinohara RT, Skoog I, Smyser CD, Sperling RA, Stein DJ, Stolicyn A, Suckling J, Sullivan G, Taki Y, Thyreau B, Toro R, Traut N, Tsvetanov KA, Turk-Browne NB, Tuulari JJ, Tzourio C, Vachon-Presseau É, Valdes-Sosa MJ, Valdes-Sosa PA, Valk SL, van Amelsvoort T, Vandekar SN, Vasung L, Victoria LW, Villeneuve S, Villringer A, Vértes PE, Wagstyl K, Wang YS, Warfield SK, Warrier V, Westman E, Westwater ML, Whalley HC, Witte AV, Yang N, Yeo B, Yun H, Zalesky A, Zar HJ, Zettergren A, Zhou JH, Ziauddeen H, Zugman A, Zuo XN; 3R-BRAIN; AIBL; Alzheimer’s Disease Neuroimaging Initiative; Alzheimer’s Disease Repository Without Borders Investigators; CALM Team; Cam-CAN; CCNP; COBRE; cVEDA; ENIGMA Developmental Brain Age Working Group; Developing Human Connectome Project; FinnBrain; Harvard Aging Brain Study; IMAGEN; KNE96; Mayo Clinic Study of Aging; NSPN; POND; PREVENT-AD Research Group; VETSA; Bullmore ET, Alexander-Bloch AF. Brain charts for the human lifespan. Nature. 2022;604(7906):525–533. doi: 10.1038/s41586-022-04554-y.

Brouwer RM, Schutte J, Janssen R, Boomsma DI, Hulshoff Pol HE, Schnack HG. The Speed of Development of Adolescent Brain Age Depends on Sex and Is Genetically Determined. Cereb Cortex. 2021;31(2):1296–1306. doi: 10.1093/cercor/bhaa296.

Couvy-Duchesne B, Faouzi J, Martin B, Thibeau-Sutre E, Wild A, Ansart M, Durrleman S, Dormont D, Burgos N, Colliot O. Ensemble Learning of Convolutional Neural Network, Support Vector Machine, and Best Linear Unbiased Predictor for Brain Age Prediction: ARAMIS Contribution to the Predictive Analytics Competition 2019 Challenge. Front Psychiatry. 2020;11:593336. doi: 10.3389/fpsyt.2020.593336.

Cole JH, Franke K. Predicting Age Using Neuroimaging: Innovative Brain Ageing Biomarkers. Trends Neurosci. 2017;40(12):681–690. doi: 10.1016/j.tins.2017.10.001.

Cole JH. Multimodality neuroimaging brain-age in UK biobank: relationship to biomedical, lifestyle, and cognitive factors. Neurobiol Aging. 2020;92:34–42. doi: 10.1016/j.neurobiolaging.2020.03.014.

de Lange AG, Cole JH. Commentary: Correction procedures in brain-age prediction. Neuroimage Clin. 2020;26:102229. doi: 10.1016/j.nicl.2020.102229..

de Lange AG, Anatürk M, Rokicki J, Han LKM, Franke K, Alnaes D, Ebmeier KP, Draganski B, Kaufmann T, Westlye LT, Hahn T, Cole JH. Mind the gap: Performance metric evaluation in brain-age prediction. Hum Brain Mapp. 2022;43(10):3113–3129. doi: 10.1002/hbm.25837.

Desikan RS, Ségonne F, Fischl B, Quinn BT, Dickerson BC, Blacker D, Buckner RL, Dale AM, Maguire RP, Hyman BT, Albert MS, Killiany RJ. An automated labeling system for subdividing the human cerebral cortex on MRI scans into gyral based regions of interest. Neuroimage. 2006;31(3):968–80. doi: 10.1016/j.neuroimage.2006.01.021.

Dima D, Modabbernia A, Papachristou E, Doucet GE, Agartz I, Aghajani M, Akudjedu TN, Albajes-Eizagirre A, Alnaes D, Alpert KI, Andersson M, Andreasen NC, Andreassen OA, Asherson P, Banaschewski T, Bargallo N, Baumeister S, Baur-Streubel R, Bertolino A, Bonvino A, Boomsma DI, Borgwardt S, Bourque J, Brandeis D, Breier A, Brodaty H, Brouwer RM, Buitelaar JK, Busatto GF, Buckner RL, Calhoun V, Canales-Rodríguez EJ, Cannon DM, Caseras X, Castellanos FX, Cervenka S, Chaim-Avancini TM, Ching CRK, Chubar V, Clark VP, Conrod P, Conzelmann A, Crespo-Facorro B, Crivello F, Crone EA, Dannlowski U, Dale AM, Davey C, de Geus EJC, de Haan L, de Zubicaray GI, den Braber A, Dickie EW, Di Giorgio A, Doan NT, Dørum ES, Ehrlich S, Erk S, Espeseth T, Fatouros-Bergman H, Fisher SE, Fouche JP, Franke B, Frodl T, Fuentes-Claramonte P, Glahn DC, Gotlib IH, Grabe HJ, Grimm O, Groenewold NA, Grotegerd D, Gruber O, Gruner P, Gur RE, Gur RC, Hahn T, Harrison BJ, Hartman CA, Hatton SN, Heinz A, Heslenfeld DJ, Hibar DP, Hickie IB, Ho BC, Hoekstra PJ, Hohmann S, Holmes AJ, Hoogman M, Hosten N, Howells FM, Hulshoff Pol HE, Huyser C, Jahanshad N, James A, Jernigan TL, Jiang J, Jönsson EG, Joska JA, Kahn R, Kalnin A, Kanai R, Klein M, Klyushnik TP, Koenders L, Koops S, Krämer B, Kuntsi J, Lagopoulos J, Lázaro L, Lebedeva I, Lee WH, Lesch KP, Lochner C, Machielsen MWJ, Maingault S, Martin NG, Martínez-Zalacaín I, Mataix-Cols D, Mazoyer B, McDonald C, McDonald BC, McIntosh AM, McMahon KL, McPhilemy G, Meinert S, Menchón JM, Medland SE, Meyer-Lindenberg A, Naaijen J, Najt P, Nakao T, Nordvik JE, Nyberg L, Oosterlaan J, de la Foz VO, Paloyelis Y, Pauli P, Pergola G, Pomarol-Clotet E, Portella MJ, Potkin SG, Radua J, Reif A, Rinker DA, Roffman JL, Rosa PGP, Sacchet MD, Sachdev PS, Salvador R, Sánchez-Juan P, Sarró S, Satterthwaite TD, Saykin AJ, Serpa MH, Schmaal L, Schnell K, Schumann G, Sim K, Smoller JW, Sommer I, Soriano-Mas C, Stein DJ, Strike LT, Swagerman SC, Tamnes CK, Temmingh HS, Thomopoulos SI, Tomyshev AS, Tordesillas-Gutiérrez D, Trollor JN, Turner JA, Uhlmann A, van den Heuvel OA, van den Meer D, van der Wee NJA, van Haren NEM, Van’t Ent D, van Erp TGM, Veer IM, Veltman DJ, Voineskos A, Völzke H, Walter H, Walton E, Wang L, Wang Y, Wassink TH, Weber B, Wen W, West JD, Westlye LT, Whalley H, Wierenga LM, Williams SCR, Wittfeld K, Wolf DH, Worker A, Wright MJ, Yang K, Yoncheva Y, Zanetti MV, Ziegler GC, Thompson PM, Frangou S; Karolinska Schizophrenia Project (KaSP). Subcortical volumes across the lifespan: Data from 18,605 healthy individuals aged 3-90 years. Hum Brain Mapp. 2022;43(1):452–469. doi: 10.1002/hbm.25320.

Elliott ML, Belsky DW, Knodt AR, Ireland D, Melzer TR, Poulton R, Ramrakha S, Caspi A, Moffitt TE, Hariri AR. Brain-age in midlife is associated with accelerated biological aging and cognitive decline in a longitudinal birth cohort. Mol Psychiatry. 2021;26(8):3829–3838. doi: 10.1038/s41380-019-0626-7.

Fischl B, Salat DH, Busa E, Albert M, Dieterich M, Haselgrove C, van der Kouwe A, Killiany R, Kennedy D, Klaveness S, Montillo A, Makris N, Rosen B, Dale AM. Whole brain segmentation: automated labeling of neuroanatomical structures in the human brain. Neuron. 2002;33(3):341–55. doi: 10.1016/s0896-6273(02)00569-x.

Frangou S, Modabbernia A, Williams SCR, Papachristou E, Doucet GE, Agartz I, Aghajani M, Akudjedu TN, Albajes-Eizagirre A, Alnaes D, Alpert KI, Andersson M, Andreasen NC, Andreassen OA, Asherson P, Banaschewski T, Bargallo N, Baumeister S, Baur-Streubel R, Bertolino A, Bonvino A, Boomsma DI, Borgwardt S, Bourque J, Brandeis D, Breier A, Brodaty H, Brouwer RM, Buitelaar JK, Busatto GF, Buckner RL, Calhoun V, Canales-Rodríguez EJ, Cannon DM, Caseras X, Castellanos FX, Cervenka S, Chaim-Avancini TM, Ching CRK, Chubar V, Clark VP, Conrod P, Conzelmann A, Crespo-Facorro B, Crivello F, Crone EA, Dale AM, Dannlowski U, Davey C, de Geus EJC, de Haan L, de Zubicaray GI, den Braber A, Dickie EW, Di Giorgio A, Doan NT, Dørum ES, Ehrlich S, Erk S, Espeseth T, Fatouros-Bergman H, Fisher SE, Fouche JP, Franke B, Frodl T, Fuentes-Claramonte P, Glahn DC, Gotlib IH, Grabe HJ, Grimm O, Groenewold NA, Grotegerd D, Gruber O, Gruner P, Gur RE, Gur RC, Hahn T, Harrison BJ, Hartman CA, Hatton SN, Heinz A, Heslenfeld DJ, Hibar DP, Hickie IB, Ho BC, Hoekstra PJ, Hohmann S, Holmes AJ, Hoogman M, Hosten N, Howells FM, Hulshoff Pol HE, Huyser C, Jahanshad N, James A, Jernigan TL, Jiang J, Jönsson EG, Joska JA, Kahn R, Kalnin A, Kanai R, Klein M, Klyushnik TP, Koenders L, Koops S, Krämer B, Kuntsi J, Lagopoulos J, Lázaro L, Lebedeva I, Lee WH, Lesch KP, Lochner C, Machielsen MWJ, Maingault S, Martin NG, Martínez-Zalacaín I, Mataix-Cols D, Mazoyer B, McDonald C, McDonald BC, McIntosh AM, McMahon KL, McPhilemy G, Meinert S, Menchón JM, Medland SE, Meyer-Lindenberg A, Naaijen J, Najt P, Nakao T, Nordvik JE, Nyberg L, Oosterlaan J, de la Foz VO, Paloyelis Y, Pauli P, Pergola G, Pomarol-Clotet E, Portella MJ, Potkin SG, Radua J, Reif A, Rinker DA, Roffman JL, Rosa PGP, Sacchet MD, Sachdev PS, Salvador R, Sánchez-Juan P, Sarró S, Satterthwaite TD, Saykin AJ, Serpa MH, Schmaal L, Schnell K, Schumann G, Sim K, Smoller JW, Sommer I, Soriano-Mas C, Stein DJ, Strike LT, Swagerman SC, Tamnes CK, Temmingh HS, Thomopoulos SI, Tomyshev AS, Tordesillas-Gutiérrez D, Trollor JN, Turner JA, Uhlmann A, van den Heuvel OA, van den Meer D, NJA van der Wee, van Haren NEM, van ‘t Ent D, van Erp TGM, Veer IM, Veltman DJ, Voineskos A, Völzke H, Walter H, Walton E, Wang L, Wang Y, Wassink TH, Weber B, Wen W, West JD, Westlye LT, Whalley H, Wierenga LM, Wittfeld K, Wolf DH, Worker A, Wright MJ, Yang K, Yoncheva Y, Zanetti MV, Ziegler GC; Karolinska Schizophrenia Project (KaSP); Thompson PM, Dima D. Cortical thickness across the lifespan: Data from 17,075 healthy individuals aged 3-90 years. Hum Brain Mapp. 2022;43(1):431–451. doi: 10.1002/hbm.25364.

Franke K, Gaser C, Manor B, Novak V. Advanced BrainAGE in older adults with type 2 diabetes mellitus. Front Aging Neurosci. 2013;5:90. doi: 10.3389/fnagi.2013.00090.

Franke K, Gaser C. Ten Years of BrainAGE as a Neuroimaging Biomarker of Brain Aging: What Insights Have We Gained? Front Neurol. 2019;10:789. doi: 10.3389/fneur.2019.00789.

Ge R, Yu, Y., Qi, Y. X., Fan, Y. V., Chen, S., Gao, C., … Frangou, S. (2023). Normative Modeling of Brain Morphometry Across the Lifespan using CentileBrain: Algorithm Benchmarking and Model Optimization. bioRxiv. doi:10.1101/2023.01.30.523509.

Goodfellow I. Bengio Y, Courville A. Deep Learning. MIT Press, 2016.

Goyal MS, Blazey TM, Su Y, Couture LE, Durbin TJ, Bateman RJ, Benzinger TL, Morris JC, Raichle ME, Vlassenko AG. Persistent metabolic youth in the aging female brain. Proc Natl Acad Sci U S A. 2019;116(8):3251–3255. doi: 10.1073/pnas.1815917116

Grinsztajn L, Oyallon E, Varoquaux G. Why do tree-based models still outperform deep learning on typical tabular data? 36^th^ Conference on Neural Information Processing Systems(NeurIPS2022),Track on Datasets and Benchmarks, 2022.

Han LKM, Dinga R, Hahn T, Ching CRK, Eyler LT, Aftanas L, Aghajani M, Aleman A, Baune BT, Berger K, Brak I, Filho GB, Carballedo A, Connolly CG, Couvy-Duchesne B, Cullen KR, Dannlowski U, Davey CG, Dima D, Duran FLS, Enneking V, Filimonova E, Frenzel S, Frodl T, Fu CHY, Godlewska BR, Gotlib IH, Grabe HJ, Groenewold NA, Grotegerd D, Gruber O, Hall GB, Harrison BJ, Hatton SN, Hermesdorf M, Hickie IB, Ho TC, Hosten N, Jansen A, Kähler C, Kircher T, Klimes-Dougan B, Krämer B, Krug A, Lagopoulos J, Leenings R, MacMaster FP, MacQueen G, McIntosh A, McLellan Q, McMahon KL, Medland SE, Mueller BA, Mwangi B, Osipov E, Portella MJ, Pozzi E, Reneman L, Repple J, Rosa PGP, Sacchet MD, Sämann PG, Schnell K, Schrantee A, Simulionyte E, Soares JC, Sommer J, Stein DJ, Steinsträter O, Strike LT, Thomopoulos SI, van Tol MJ, Veer IM, Vermeiren RRJM, Walter H, van der Wee NJA, van der Werff SJA, Whalley H, Winter NR, Wittfeld K, Wright MJ, Wu MJ, Völzke H, Yang TT, Zannias V, de Zubicaray GI, Zunta-Soares GB, Abé C, Alda M, Andreassen OA, Bøen E, Bonnin CM, Canales-Rodriguez EJ, Cannon D, Caseras X, Chaim-Avancini TM, Elvsåshagen T, Favre P, Foley SF, Fullerton JM, Goikolea JM, Haarman BCM, Hajek T, Henry C, Houenou J, Howells FM, Ingvar M, Kuplicki R, Lafer B, Landén M, Machado-Vieira R, Malt UF, McDonald C, Mitchell PB, Nabulsi L, Otaduy MCG, Overs BJ, Polosan M, Pomarol-Clotet E, Radua J, Rive MM, Roberts G, Ruhe HG, Salvador R, Sarró S, Satterthwaite TD, Savitz J, Schene AH, Schofield PR, Serpa MH, Sim K, Soeiro-de-Souza MG, Sutherland AN, Temmingh HS, Timmons GM, Uhlmann A, Vieta E, Wolf DH, Zanetti MV, Jahanshad N, Thompson PM, Veltman DJ, Penninx BWJH, Marquand AF, Cole JH, Schmaal L. Brain aging in major depressive disorder: results from the ENIGMA major depressive disorder working group. Mol Psychiatry. 2021;26(9):5124–5139. doi: 10.1038/s41380-020-0754-0.

He T, Kong R, Holmes AJ, Nguyen M, Sabuncu MR, Eickhoff SB, Bzdok D, Feng J, Yeo BTT. Deep neural networks and kernel regression achieve comparable accuracies for functional connectivity prediction of behavior and demographics. Neuroimage. 2020;206:116276. doi: 10.1016/j.neuroimage.2019.116276..

Hogstrom LJ, Westlye LT, Walhovd KB, Fjell AM. The structure of the cerebral cortex across adult life: age-related patterns of surface area, thickness, and gyrification. Cereb Cortex. 2013;23(11):2521–30. doi: 10.1093/cercor/bhs231.

Liang H, Zhang F, Niu X. Investigating systematic bias in brain age estimation with application to post-traumatic stress disorders. Hum Brain Mapp. 2019;40(11):3143–3152. doi: 10.1002/hbm.24588.

Lombardi A, Amoroso N, Diacono D, Monaco A, Tangaro S, Bellotti R. Extensive Evaluation of Morphological Statistical Harmonization for Brain Age Prediction. Brain Sci. 2020;10(6):364. doi: 10.3390/brainsci10060364.

Luna A, Bernanke J, Kim K, Aw N, Dworkin JD, Cha J, Posner J. Maturity of gray matter structures and white matter connectomes, and their relationship with psychiatric symptoms in youth. Hum Brain Mapp. 2021;42(14):4568–4579. doi: 10.1002/hbm.25565.

Lund MJ, Alnæs D, de Lange AG, Andreassen OA, Westlye LT, Kaufmann T. Brain age prediction using fMRI network coupling in youths and associations with psychiatric symptoms. Neuroimage Clin. 2022;33:102921.

Modabbernia A, Whalley HC, Glahn DC, Thompson PM, Kahn RS, Frangou S. Systematic evaluation of machine learning algorithms for neuroanatomically-based age prediction in youth. Hum Brain Mapp. 2022;43(17):5126–5140. doi: 10.1002/hbm.26010.

More S, Antonopoulos G, Hoffstaedter F, Caspers J, Eickhoff SB, Patil KR; Alzheimer’s Disease Neuroimaging Initiative. Brain-age prediction: A systematic comparison of machine learning workflows. Neuroimage. 2023;270:119947. doi: 10.1016/j.neuroimage.2023.119947.

Niu X, Zhang F, Kounios J, Liang H. Improved prediction of brain age using multimodal neuroimaging data. Hum Brain Mapp. 2020;41(6):1626–1643. doi: 10.1002/hbm.24899.

Pomponio R, Erus G, Habes M, Doshi J, Srinivasan D, Mamourian E, Bashyam V, Nasrallah IM, Satterthwaite TD, Fan Y, Launer LJ, Masters CL, Maruff P, Zhuo C, Völzke H, Johnson SC, Fripp J, Koutsouleris N, Wolf DH, Gur R, Gur R, Morris J, Albert MS, Grabe HJ, Resnick SM, Bryan RN, Wolk DA, Shinohara RT, Shou H, Davatzikos C. Harmonization of large MRI datasets for the analysis of brain imaging patterns throughout the lifespan. Neuroimage. 2020;208:116450. doi: 10.1016/j.neuroimage.2019.116450.

Rokicki J, Wolfers T, Nordhøy W, Tesli N, Quintana DS, Alnaes D, Richard G, de Lange AG, Lund MJ, Norbom L, Agartz I, Melle I, Naerland T, Selbaek G, Persson K, Nordvik JE, Schwarz E, Andreassen OA, Kaufmann T, Westlye LT. Multimodal imaging improves brain age prediction and reveals distinct abnormalities in patients with psychiatric and neurological disorders. Hum Brain Mapp. 2021;42(6):1714–1726. doi: 10.1002/hbm.25323

Schölkopf B, Smola AJ. Learning with Kernels: Support Vector Machines, Regularization, Optimization, and Beyond. MIT Press, 2002.

Schulz MA, Yeo BTT, Vogelstein JT, Mourao-Miranada J, Kather JN, Kording K, Richards B, Bzdok D. Different scaling of linear models and deep learning in UKBiobank brain images versus machine-learning datasets. Nat Commun. 2020 25;11(1):4238. doi: 10.1038/s41467-020-18037-z.

Valizadeh SA, Hänggi J, Mérillat S, Jäncke L. Age prediction on the basis of brain anatomical measures. Hum Brain Mapp. 2017;38(2):997–1008. doi: 10.1002/hbm.23434.

Zhou Z, Li H, Srinivasan D, Abdulkadir A, Nasrallah IM, Wen J, Doshi J, Erus G, Mamourian E, Bryan NR, Wolk DA, Beason-Held L, Resnick SM, Satterthwaite TD, Davatzikos C, Shou H, Fan Y; ISTAGING Consortium. Multiscale functional connectivity patterns of the aging brain learned from harmonized rsfMRI data of the multi-cohort iSTAGING study. Neuroimage. 2023;269:119911. doi: 10.1016/j.neuroimage.2023.119911.

