## supplementary text for "Brain-Age Prediction: Systematic Evaluation of Site Effects, and Sample Age Range and Size"

**Table of Contents**

**S1. Age-by-sex distribution of the study sample**
Supplementary Figure S1. Distribution of participants’ chronological age
Supplementary Table S1. Details of the discovery sample
Supplementary Table S2. Details of the independent replication and consistency samples
**S2. Neuroimaging data quality assessment**
**S3. Effect of site handling methods for the sex-specific models**
Supplementary Figure S2. Performance metrics derived from repeated cross-validation in the discovery sample across different age range partition options in females
Supplementary Figure S3. Performance metrics derived from repeated cross-validation in the discovery sample across different age range partition options in males
Supplementary Table S3. Females Only. Mean and standard deviation (SD) of the mean absolute error (MAE) and of the correlation coefficient between the chronological age and brain-age in the discovery sample
Supplementary Table S4. Males Only. Mean and standard deviation (SD) of the mean absolute error (MAE) and of the correlation coefficient between the chronological age and brain-age.in the discovery sample
Supplementary Table S5. Females Only. Mean and standard deviation (SD) of the mean absolute error (MAE) and of the correlation coefficient between the chronological age and brain-age.in the replication sample
Supplementary Table S6. Males Only. Mean and standard deviation (SD) of the mean absolute error (MAE) and of the correlation coefficient between the chronological age and brain-age.in the replication sample
**S4. Effect of age range for the sex-specific models**
Supplementary Figure S4. Performance metrics derived from the application of the models pre-trained on the discovery sample to the different age bins of the replication sample
Supplementary Figure S5. Scatterplots of the relationship between brain age and chronological age in different age bins
**S5. Effect of sample size for the sex-specific models**
Supplementary Figure S6. Model performance as a function of sample size for the preferred model
**S6. Longitudinal consistency**
Supplementary Figure S7. Females only. Model performance on longitudinal data
Supplementary Figure S8. Males only. Model performance on longitudinal data
Supplementary Table S7. Females Only. Pre-trained model performances in the independent longitudinal-consistency sample
Supplementary Table S8. Males Only. Pre-trained model performances in the independent longitudinal-consistency sample
**S7. References**

### S1. Age-by-sex distribution of the study sample

**Figure S1. The distribution of participants' chronological age in the discovery, replication, and consistency samples**. For the discovery and replication samples, the horizontal axis depicts chronological age in years in age-bins of 10 years, while the vertical axis shows the number of participants. For the consistency sample, the horizontal axis depicts chronological age in years in age-bins of 1.5 years; and the vertical axis shows the number of participants; the age distribution of the first (blue) and second (orange) scans is depicted using distinct colors. Detailed information about the datasets is presented in Table S1 and Table S2.

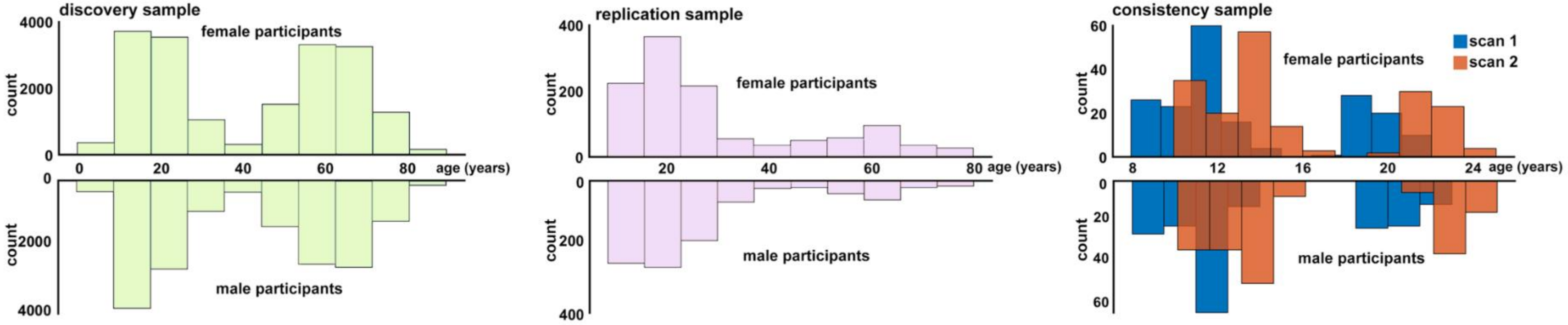

### S2. Neuroimaging data quality assessment

For samples drawn from the ENIGMA Lifespan Working Group (supplemental material Table S1), quality assessment followed the ENIGMA consortium pipeline (http://enigma.ini.usc.edu/protocols/imaging-protocols/). This pipeline was applied to 10,476 individuals (54.17% female). Data from the Adolescent Brain Cognitive Development study (N=3,116; 53.05% female) were considered high-quality based on their own quality assessment protocols (Txt file=FreeSurfer QC; item=fsqc_qc; scan excluded if 0) (Casey et al., 2018). For the remainder of the sample involving 22,091 participants (53.39% female) T1-weighted images were downloaded and segmented locally, and the results were quality assessed using Qoala-T tool (Klapwijk et al., 2019).

### S3. Effect of site handling methods in the sex-specific models

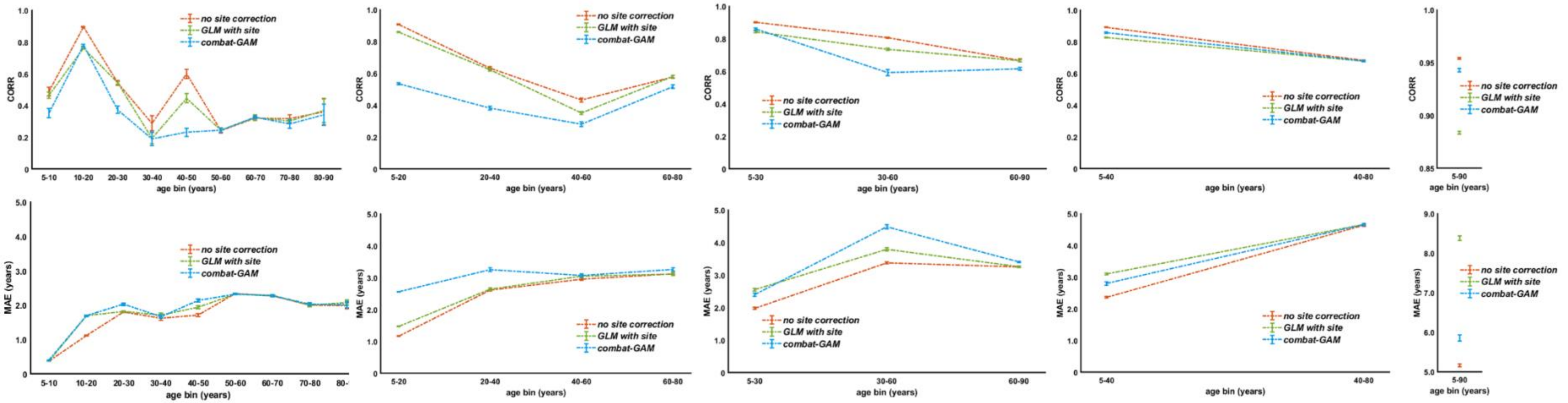


**Figure S2. Performance metrics derived from repeated cross-validation in the discovery sample across different age range partition options in females**. Each line represents one of the three site handling methods: Red=no site correction; Blue=site harmonisation with Combat-GAM; Green= site data residualization using a generalized linear model (GLM); CORR, correlation coefficient between brain-age and chronological age; MAE: mean absolute error between brain-age and chronological age.

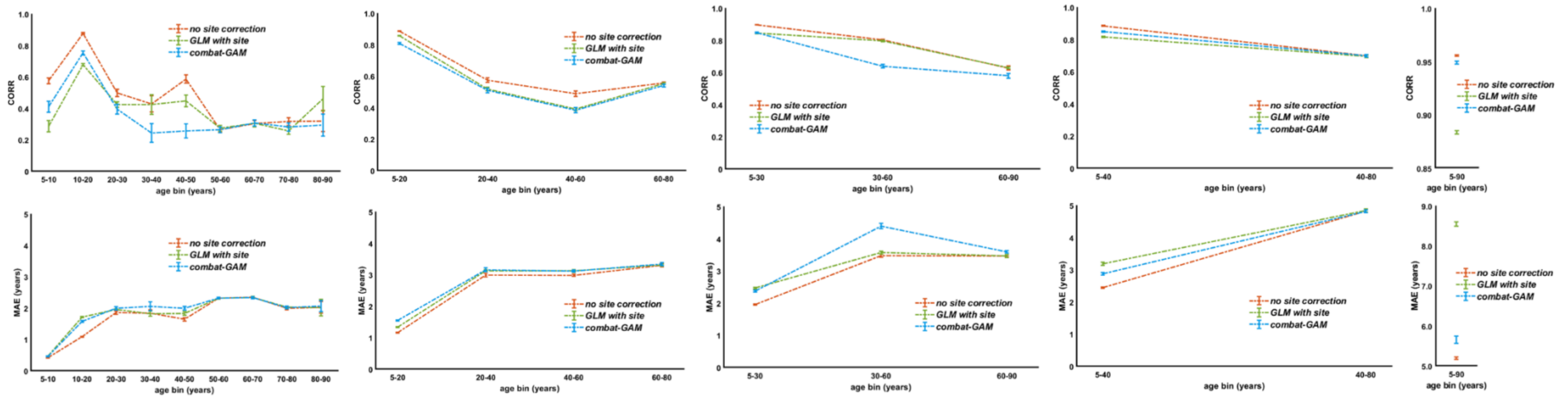


**Figure S3. Performance metrics derived from repeated cross-validation in the discovery sample across different age range partition options in males.** Each line represents one of the three site handling methods: Red=no site correction; Blue=site harmonisation with Combat-GAM; Green= site data residualization using a generalized linear model (GLM); CORR, correlation coefficient between brain-age and chronological age; MAE: mean absolute error between brain-age and chronological age.

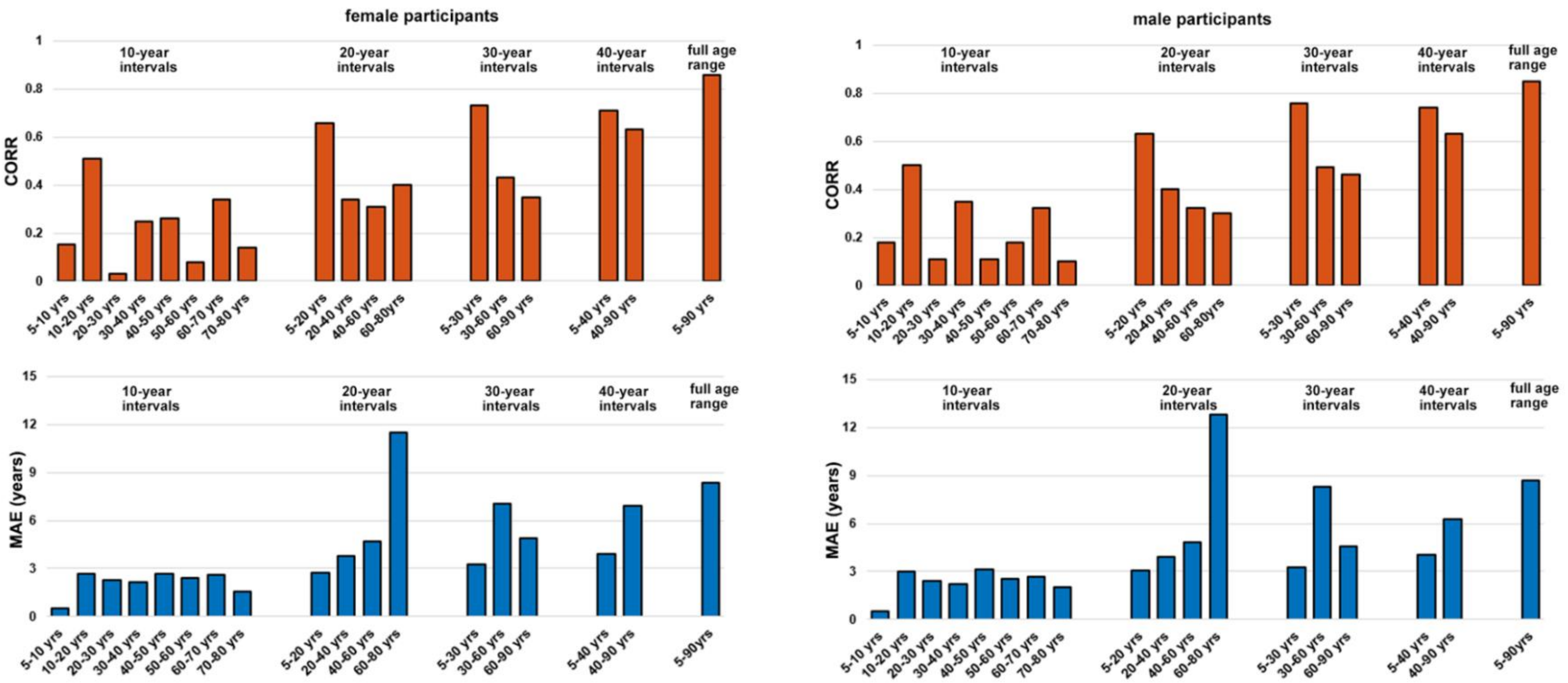


**Figure S4.** Performance metrics derived from the application of the models pre-trained on different age bins of the discovery sample to the corresponding age bins of the replication sample. The pre-trained model generated in each age bin of the discovery sample was applied to the corresponding age bin of the replication sample. The results for female participants and male participants are presented in the left panel and right panel, respectively. For female participants, average CORR values for each age bin were: 0.22 (10-year intervals), 0.43 (20-year intervals), 0.50 (30-year intervals), 0.67 (40-year intervals), and 0.86 (full age range); average MAE values for each age range were 2.10 years (10-year intervals), 5.68 years (20-year intervals), 5.03 years (30-year intervals), 5.43 years (40-year intervals), and 8.33 years (full age range). For male participants, average CORR values for each age range were 0.23 (10-year intervals), 0.41 (20-year intervals), 0.57 (30-year intervals), 0.69 (40-year intervals), and 0.85 (full age range); average MAE values for each age range were 2.32 years (10-year intervals), 6.15 years (20-year intervals), 5.37 years (30-year intervals), 5.13 years (40-year intervals), and 8.71 years (full age range).

CORR, correlation coefficient between brain-age and chronological age; MAE: mean absolute error between brain-age and chronological age.

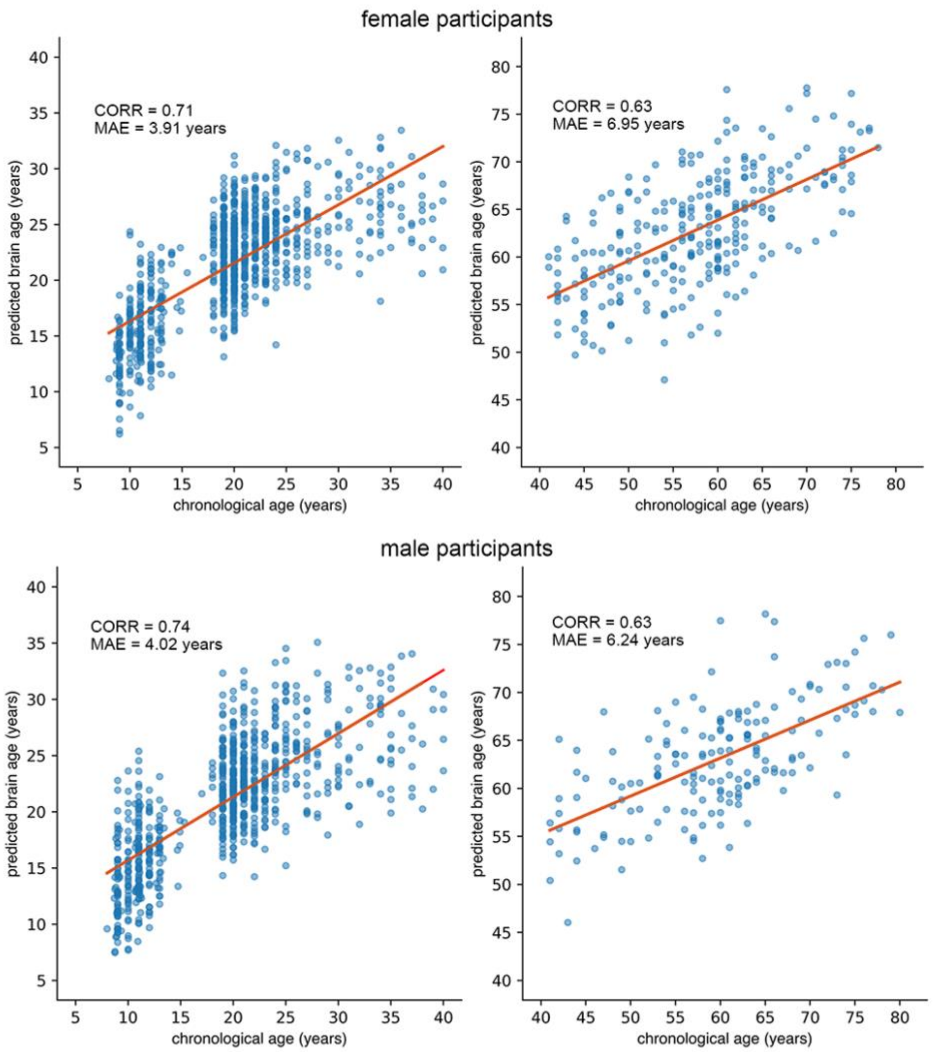


**Figure S5. Scatterplots of the relationship between brain-age and chronological age for each age bin**. The results for female participants and male participants are presented in the upper panel and lower panel, respectively. CORR, correlation coefficient between brain-age and chronological age; MAE: mean absolute error between brain-age and chronological age.

**S5. Effect of sample size for sex-specific models**

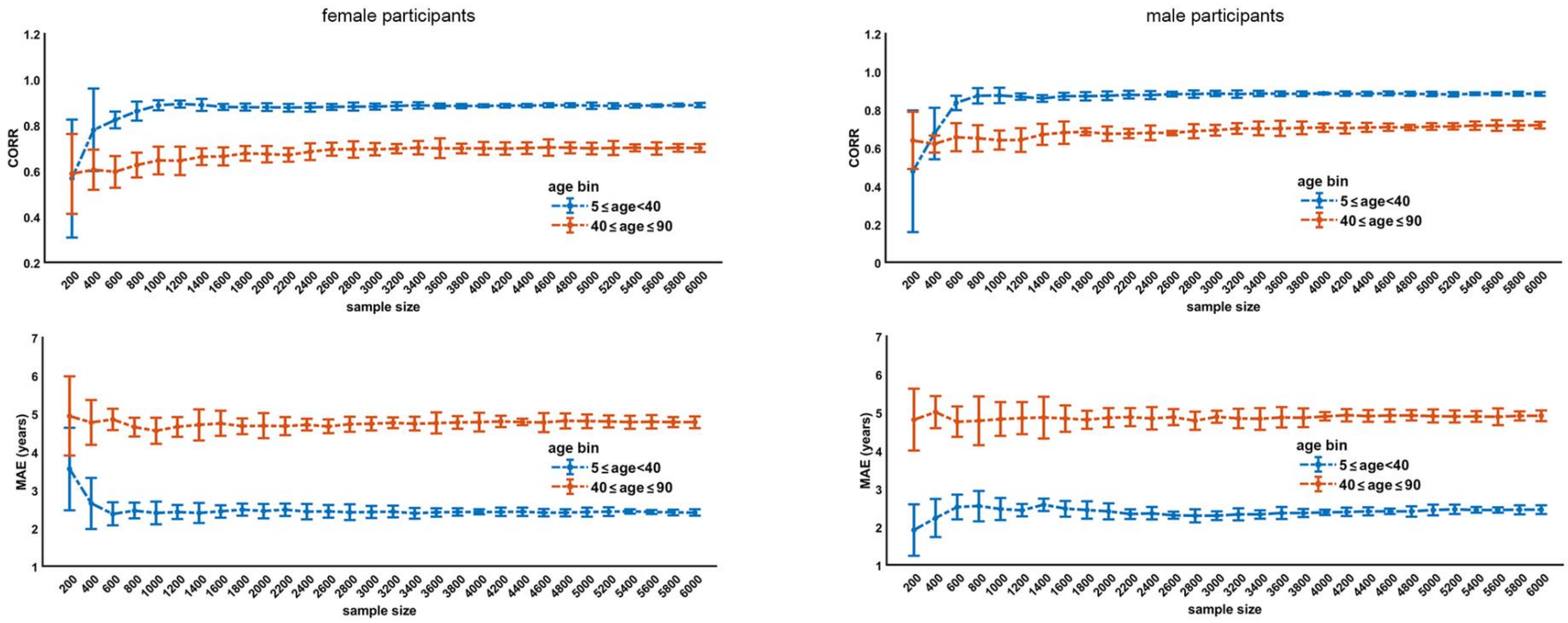


**Figure S6.** Model performance as a function of sample size in the model discovery sample with two chronological age ranges (5-40 years and 40-90 years). Model parameters for each age range were obtained by randomly resampling the model discovery sample without replacement generating subsets of 200-6,000 participants. The results for female participants and male participants are presented in the left panel and right panel, respectively.

CORR, correlation coefficient between brain-age and chronological age; MAE: mean absolute error between brain-age and chronological age.

### S6. Longitudinal consistency

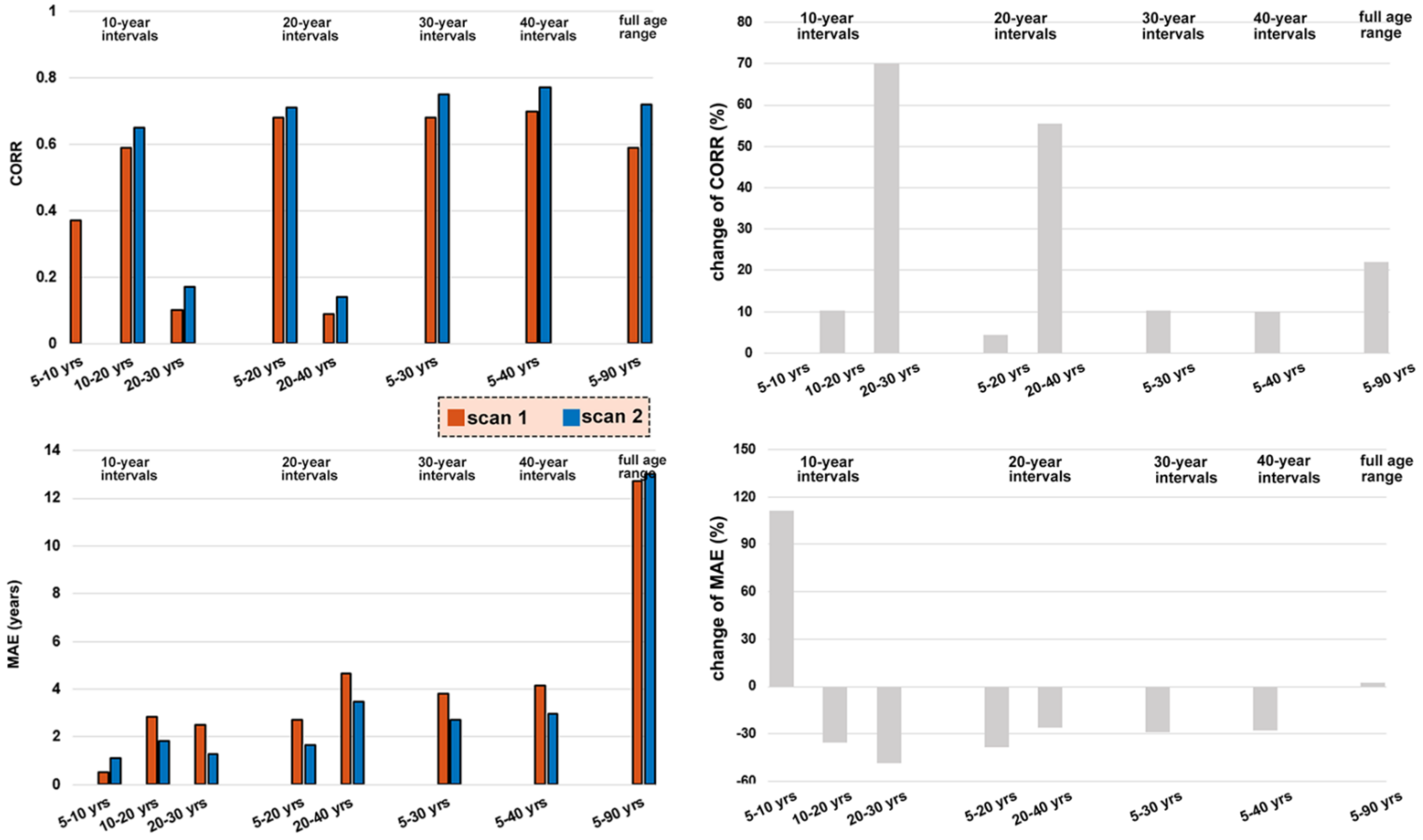


**Figure S7.** Model performance on longitudinal data of the female participants. The left panel presents the CORR and MAE values for the first and second MRI scans, while the right panel exhibits the percentage changes (%) in CORR and MAE for the second scan compared to the first scan. The results were generated by employing models that had been trained on discovery samples from each age range division and then applied to the longitudinal sample. Female and male participants were presented separately.

CORR: correlation coefficient between brain-age and chronological age; MAE: mean absolute error between brain-age and chronological age.

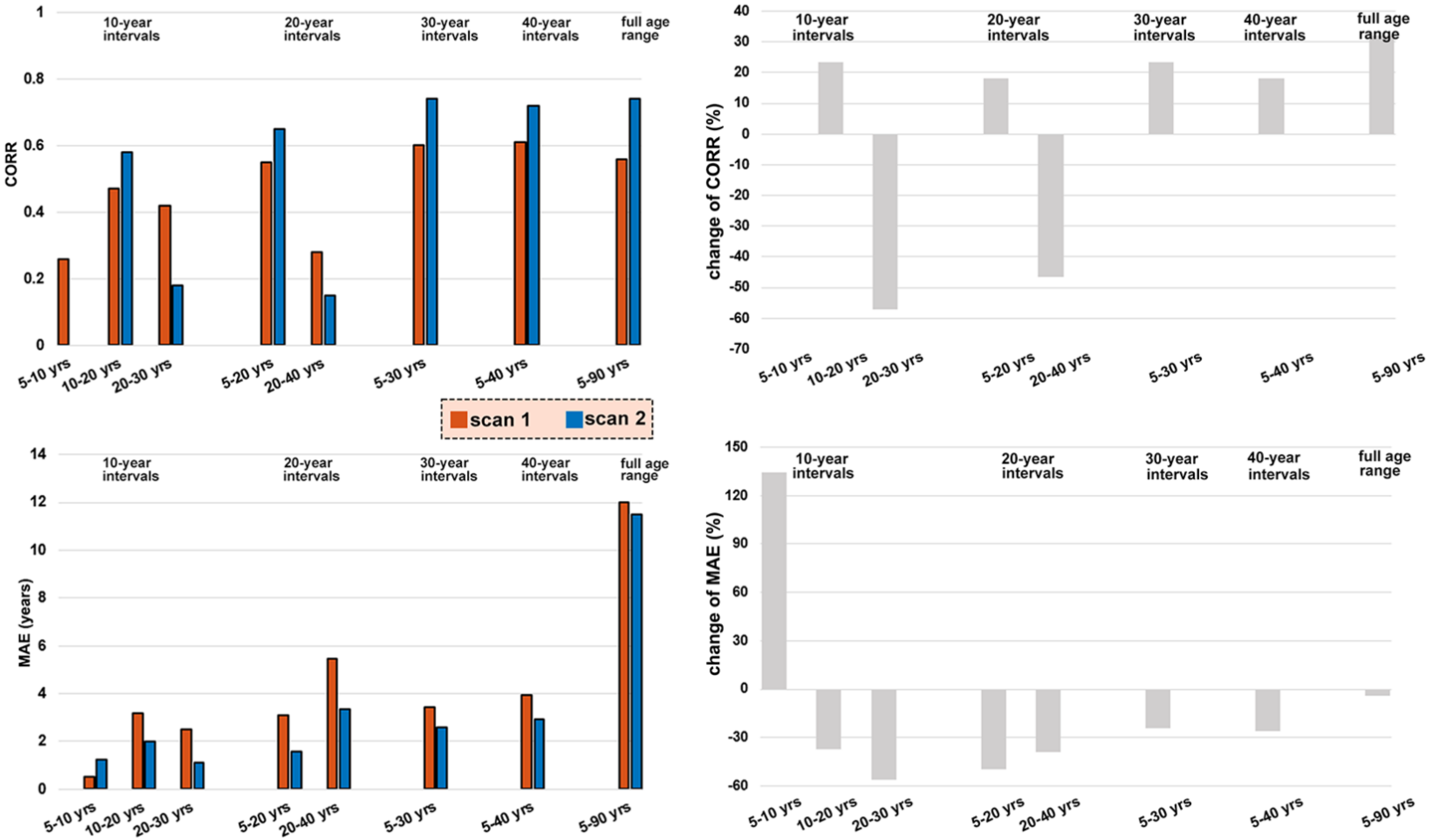


**Figure S8.** Model performance in longitudinal data of the male participants. The left panel presents the CORR and MAE values for the first and second MRI scans, while the right panel exhibits the percentage changes (%) in CORR and MAE for the second scan compared to the first scan. The results were generated by employing models that had been trained on discovery samples from each age range division and then applied to the longitudinal sample. Female and male participants were presented separately.

CORR: correlation coefficient between brain-age and chronological age; MAE: mean absolute error between brain-age and chronological age.

### S7. References


Casey, B. J., et al. (2018). The adolescent brain cognitive development (ABCD) study: imaging acquisition across 21 sites. *Developmental Cognitive Neuroscience, 32*, 43-54. doi:https://doi.org/10.1016/j.dcn.2018.03.001

Klapwijk, E. T., et al. (2019). Qoala-T: A supervised-learning tool for quality control of FreeSurfer segmented MRI data. *NeuroImage, 189*, 116-129. doi:https://doi.org/10.1016/j.neuroimage.2019.01.014
