## supplementary table 1 for "Brain-Age Prediction: Systematic Evaluation of Site Effects, and Sample Age Range and Size"

| **Supplementary Table S1. Details of the Discovery Sample** | | | | | | | |
| --- | --- | --- | --- | --- | --- | --- | --- |
| **Dataset Abbreviation** | **Country of Scanning Site** | **Scanner Strength (Tesla)** | **N** | **Age (years) Mean (SD)** | **Age Range (years)** | **Male/Female** | **Further Information and Access Requests** |
| ABCD | USA | 3T | 3116 | 9.91(0.63) | 8-11 | 1463/1653 | Access request: https://nda.nih.gov/abcd/ |
| ABIDE | USA, Germany, The Netherlands, Ireland, Belgium | 3T | 438 | 13.62(3.41) | 6-21 | 355/83 | Access request: http://fcon_1000.projects.nitrc.org/indi/abide/abide_I.html |
| ABIDE II | USA, The Netherlands, Switzerland, France, Belgium, Ireland | 1.5T;3T | 433 | 11.52(3.41) | 5-21 | 292/141 | Access request: http://fcon_1000.projects.nitrc.org/indi/abide/abide_II.html |
| ADHD200 | USA, The Netherlands, China | 1.5T; 3T | 389 | 12.12(3.32) | 7-21 | 181/208 | Access request: http://fcon_1000.projects.nitrc.org/indi/adhd200/index.html |
| AMC | The Netherlands | 3T | 99 | 22.89(3.39) | 17-32 | 65/34 | https://www.ncbi.nlm.nih.gov/pmc/articles/PMC8675431/ https://www.ncbi.nlm.nih.gov/pmc/articles/PMC8675429/ Access request to PI |
| Ann Arbor, a | USA | 3T | 25 | 21(7.42) | 13-41 | 22/3 | Access request: http://fcon_1000.projects.nitrc.org/fcpClassic/FcpTable.html |
| Ann Arbor, b | USA | 3T | 35 | 47.08(26.14) | 19-80 | 16/19 | Access request: http://fcon_1000.projects.nitrc.org/fcpClassic/FcpTable.html |
| Atlanta | USA | 3T | 28 | 30.89(9.89) | 22-57 | 13/15 | Access request: http://fcon_1000.projects.nitrc.org/fcpClassic/FcpTable.html |
| Baltimore | USA | 3T | 23 | 29.26(5.46) | 20–40 | 8/15 | Access request: http://fcon_1000.projects.nitrc.org/fcpClassic/FcpTable.html |
| Bangor | UK | 3T | 20 | 23.4(5.32) | 19-38 | 20/0 | Access request: http://fcon_1000.projects.nitrc.org/fcpClassic/FcpTable.html |
| Barcelona 3T | Spain | 3T | 44 | 14.57(2.1) | 11-17 | 24/20 | https://www.ncbi.nlm.nih.gov/pmc/articles/PMC8675431/ https://www.ncbi.nlm.nih.gov/pmc/articles/PMC8675429/ Access request to PI |
| Beijing_Zang | China | 3T | 198 | 21.16(1.83) | 18-26 | 76/122 | Access request: http://fcon_1000.projects.nitrc.org/fcpClassic/FcpTable.html |
| Berlin_Margulies | Germany | 3T | 26 | 29.77(5.21) | 23–44 | 13/13 | Access request: http://fcon_1000.projects.nitrc.org/fcpClassic/FcpTable.html |
| Betula | Sweden | 3T | 298 | 62.89(13.09) | 25-82 | 144/154 | Access request: https://www.umu.se/en/research/projects/betula---aging-memory-and-dementia/ |
| BIG | The Netherlands | 1.5T;3T | 1301 | 24.04(8.06) | 18-71 | 556/745 | Access request: https://www.ru.nl/donders/research/research-facilities-projects/cognomics/big-project/big-database/ |
| BIL&GIN | France | 3T | 452 | 26.75(7.73) | 18-57 | 220/232 | Access request: https://www.gin.cnrs.fr/en/current-research/axis2/bilgin-en/ |
| Bonn | Germany | 3T | 174 | 38.84(6.49) | 29-50 | 174/0 | https://www.ncbi.nlm.nih.gov/pmc/articles/PMC8675431/ https://www.ncbi.nlm.nih.gov/pmc/articles/PMC8675429/ Access request to PI |
| BrainSCALE | The Netherlands | 1.5T | 277 | 10.02(1.38) | 9-15 | 131/146 | https://www.ncbi.nlm.nih.gov/pmc/articles/PMC8675431/ https://www.ncbi.nlm.nih.gov/pmc/articles/PMC8675429/ Access request to PI |
| BRCATLAS | England | 3T | 163 | 39.66(17.17) | 18-84 | 84/79 | https://www.ncbi.nlm.nih.gov/pmc/articles/PMC8675431/ https://www.ncbi.nlm.nih.gov/pmc/articles/PMC8675429/ Access request to PI |
| Cambridge | USA | 3T | 198 | 21.03 (2.31) | 18-30 | 75/123 | Access request: http://fcon_1000.projects.nitrc.org/fcpClassic/FcpTable.html |
| Cam-CAN | UK | 3T | 643 | 54.56(18.55) | 18-89 | 314/329 | Access request: https://camcan-archive.mrc-cbu.cam.ac.uk/dataaccess/ |
| Cardiff | Wales | 3T | 318 | 25.29(7.4) | 18-58 | 89/229 | https://www.ncbi.nlm.nih.gov/pmc/articles/PMC8675431/ https://www.ncbi.nlm.nih.gov/pmc/articles/PMC8675429/ Access request to PI |
| CLiNG | Germany | 3T | 323 | 25.18(5.28) | 18-58 | 132/191 | https://www.ncbi.nlm.nih.gov/pmc/articles/PMC8675431/ https://www.ncbi.nlm.nih.gov/pmc/articles/PMC8675429/ Access request to PI |
| CODE | Germany | 3T | 74 | 39.99(13.32) | 20-64 | 31/43 | Access request: https://www.ncbi.nlm.nih.gov/pmc/articles/PMC4651841/ |
| COMPULS/TS EUROTRAIN | The Netherlands | 3T | 54 | 10.86(1.02) | 8-13 | 37/17 | [https://www.ncbi.nlm.nih.gov/pmc/articles/PMC4994475/ Access request to PI](https://www.ncbi.nlm.nih.gov/pmc/articles/PMC4994475/Access%20request%20to%20PI) |
| GSP | USA | 3T | 2010 | 27.23(16.53) | 18-90 | 895/1115 | Access request: https://dataverse.harvard.edu/dataset.xhtml?persistentId=doi:10.7910/DVN/25833 |
| HCP Adult Brain | USA | 3T | 1113 | 28.8(3.7) | 22-37 | 507/606 | <https://www.humanconnectome.org/study/hcp-young-adult/overview> |
| HCP Aging | USA | 3T | 725 | 60.36(15.73) | 36-100 | 319/406 | <https://www.humanconnectome.org/study/hcp-lifespan-aging> |
| HCP Development | USA | 3T | 652 | 14.44(4.06) | 5-22 | 301/351 | <https://www.humanconnectome.org/study/hcp-lifespan-development/overview> |
| Healthy Brain Network/CMI | USA | 1.5T;3T | 214 | 9.97(3.4) | 5-21 | 118/96 | Access request: http://fcon_1000.projects.nitrc.org/indi/cmi_healthy_brain_network/ |
| ICBM | Canada | 3T; 1.5T; 2T | 86 | 44.19(17.92) | 19-85 | 41/45 | Access request: http://fcon_1000.projects.nitrc.org/fcpClassic/FcpTable.html |
| IMAGEN | England, Ireland, France, Germany | 3T | 1840 | 14.45(0.4) | 13-16 | 903/937 | https://imagen-europe.com/ |
| IMH | Singapore | 3T | 79 | 31.99(10.03) | 20-59 | 50/29 | https://www.ncbi.nlm.nih.gov/pmc/articles/PMC8675431/ https://www.ncbi.nlm.nih.gov/pmc/articles/PMC8675429/ Access request to PI |
| Indiana | USA | 3T | 201 | 27.54(20.02) | 6-87 | 97/104 | https://www.ncbi.nlm.nih.gov/pmc/articles/PMC8675431/ https://www.ncbi.nlm.nih.gov/pmc/articles/PMC8675429/ Access request to PI |
| IRCCS-FSL | Italy | 3T | 148 | 66.9(5.31) | 60-83 | 61/87 | https://www.ncbi.nlm.nih.gov/pmc/articles/PMC8675431/ https://www.ncbi.nlm.nih.gov/pmc/articles/PMC8675429/ Access request to PI |
| Leiden | The Netherlands | 3T | 576 | 16.84(4.79) | 8-29 | 281/295 | https://www.ncbi.nlm.nih.gov/pmc/articles/PMC8675431/ https://www.ncbi.nlm.nih.gov/pmc/articles/PMC8675429/ Access request to PI |
| Leiden_2180 | The Netherlands | 3T | 12 | 23.00(2.49) | 20-27 | 12/0 | Access request: http://fcon_1000.projects.nitrc.org/fcpClassic/FcpTable.html |
| Leiden_2200 | The Netherlands | 3T | 19 | 21.69(2.56) | 18-28 | 11/8 | Access request: http://fcon_1000.projects.nitrc.org/fcpClassic/FcpTable.html |
| MAS | Australia | 3T | 532 | 78.41(4.68) | 70-91 | 242/290 | [https://cheba.unsw.edu.au/research-projects/sydney-memory-and-ageing-study Access request to PI](https://cheba.unsw.edu.au/research-projects/sydney-memory-and-ageing-studyAccess%20request%20to%20PI) |
| MCIC | USA | 1.5T; 3T | 93 | 32.67(12.03) | 18-60 | 63/30 | [Access request: https://www.nitrc.org/projects/mcic/](https://www.ncbi.nlm.nih.gov/pmc/articles/PMC3727653/Access%20request%20to%20PI) |
| Melbourne | Australia | 3T | 102 | 19.58(2.98) | 15-26 | 48/54 | https://www.ncbi.nlm.nih.gov/pmc/articles/PMC8675431/ https://www.ncbi.nlm.nih.gov/pmc/articles/PMC8675429/ Access request to PI |
| Milwaukee_b | USA | 3T | 46 | 53.59(5.8) | 44-65 | 15/31 | Access request: http://fcon_1000.projects.nitrc.org/fcpClassic/FcpTable.html |
| Muenster | Germany | 3T | 752 | 35.2(12.07) | 17-65 | 328/424 | https://www.ncbi.nlm.nih.gov/pmc/articles/PMC8675431/ https://www.ncbi.nlm.nih.gov/pmc/articles/PMC8675429/ Access request to PI |
| Munchen | Germany | 3T | 16 | 68.44(3.97) | 63-74 | 10/6 | Access request: http://fcon_1000.projects.nitrc.org/fcpClassic/FcpTable.html |
| Neuroventure | Canada | 3T | 137 | 13.66(0.64) | 12-15 | 62/75 | https://www.ncbi.nlm.nih.gov/pmc/articles/PMC8675431/ https://www.ncbi.nlm.nih.gov/pmc/articles/PMC8675429/ Access request to PI |
| Newark | USA | 3T | 19 | 24.11(3.91) | 21-39 | 9/10 | Access request: http://fcon_1000.projects.nitrc.org/fcpClassic/FcpTable.html |
| NewYork_b | USA | 3T | 20 | 29.75(9.94) | 18-46 | 8/12 | Access request: http://fcon_1000.projects.nitrc.org/fcpClassic/FcpTable.html |
| NUIG | Ireland | 1.5T | 93 | 36.14(11.55) | 18-58 | 54/39 | https://www.ncbi.nlm.nih.gov/pmc/articles/PMC8675431/ https://www.ncbi.nlm.nih.gov/pmc/articles/PMC8675429/ Access request to PI |
| NYU_TRT | USA | 3T | 25 | 29.44(8.64) | 22-49 | 10/15 | Access request: http://fcon_1000.projects.nitrc.org/fcpClassic/FcpTable.html |
| Olin | USA | 3T | 607 | 36.04(12.99) | 21-87 | 242/365 | https://www.ncbi.nlm.nih.gov/pmc/articles/PMC8675431/ https://www.ncbi.nlm.nih.gov/pmc/articles/PMC8675429/ Access request to PI |
| Orangeburg | USA | 1.5T | 20 | 40.65(11.03) | 25-55 | 15/5 | Access request: http://fcon_1000.projects.nitrc.org/fcpClassic/FcpTable.html |
| Oulu | Finland | 1.5T | 103 | 21.52(0.57) | 20-23 | 37/66 | Access request: http://fcon_1000.projects.nitrc.org/fcpClassic/FcpTable.html |
| Oxford | UK | 3T | 22 | 29(3.79) | 20-35 | 12/10 | Access request: http://fcon_1000.projects.nitrc.org/fcpClassic/FcpTable.html |
| PaloAlto | USA | 3T | 17 | 32.47(8.12) | 23-39 | 2/15 | Access request: http://fcon_1000.projects.nitrc.org/fcpClassic/FcpTable.html |
| PING | USA | 3T | 518 | 11.86(4.91) | 5-21 | 271/247 | http://pingstudy.ucsd.edu/welcome.html |
| QTIM | Australia | 4T | 342 | 22.64(3.35) | 16-30 | 112/230 | Access request: https://openneuro.org/datasets/ds004169/versions/1.0.6 |
| Queensland | Australia | 3T | 19 | 25.95(3.88) | 23-34 | 11/8 | Access request: http://fcon_1000.projects.nitrc.org/fcpClassic/FcpTable.html |
| Rockland | USA | 3T | 140 | 14.06(4.86) | 6-21 | 63/77 | Access request: https://fcon_1000.projects.nitrc.org/indi/pro/nki.html |
| Saint-Louis | USA | 3T | 31 | 25.1(2.31) | 21-29 | 14/17 | Access request: http://fcon_1000.projects.nitrc.org/fcpClassic/FcpTable.html |
| Sydney | Australia | 3T | 157 | 39.13(22.09) | 12-84 | 65/92 | https://www.ncbi.nlm.nih.gov/pmc/articles/PMC8675431/ https://www.ncbi.nlm.nih.gov/pmc/articles/PMC8675429/ Access request to PI |
| UK BIOBANK | UK | 3T | 14496 | 62.16(7.57) | 45-82 | 6494/8002 | [Access request: https://www.ukbiobank.ac.uk/](https://www.ukbiobank.ac.uk/) |
| UNIBA | Italy | 3T | 131 | 27.34(9.06) | 18-63 | 67/64 | https://www.ncbi.nlm.nih.gov/pmc/articles/PMC8675431/ https://www.ncbi.nlm.nih.gov/pmc/articles/PMC8675429/ Access request to PI |
| UPENN | USA | 3T | 190 | 36.05(14.04) | 16-85 | 87/103 | https://www.ncbi.nlm.nih.gov/pmc/articles/PMC8675431/ https://www.ncbi.nlm.nih.gov/pmc/articles/PMC8675429/ Access request to PI |
| UTH | USA | 1.5T; 3T | 231 | 34.4(12.63) | 18-65 | 89/142 | https://www.ncbi.nlm.nih.gov/pmc/articles/PMC8675431/ https://www.ncbi.nlm.nih.gov/pmc/articles/PMC8675429/ Access request to PI |
| Total |  |  | 35683 | 30.35(7.80) | 5-90 | 16561/19122 |  |
| ABCD Healthy, The Adolescent Brain Cognitive Development Study; ABIDE, Autism Brain Imaging Data Exchange; ADHD200, Attention Deficit Hyperactivity Disorder – ADHD200 sample; AMC, Amsterdam Medisch Centrum; Barcelona 3T, University of Barcelona; Betula, The Betula Project, Swedish longitudinal study on aging, memory, and dementia; BIG, Brain Imaging Genetics; BIL&GIN, Brain Imaging of Lateralization by the Groupe d'Imagerie Neurofonctionnelle; Bonn, University of Bonn; BrainSCALE, Brain Structure and Cognition: an adolescence longitudinal twin study; BRCATLAS, Biomedical Research Centre Atlas; Cam-CAN, Cambridge Centre for Ageing and Neuroscience; Cardiff, Cardiff University; CLiNG, Clinical Neuroscience Göttingen; CODE, formerly Cognitive Behavioral Analysis System of Psychotherapy (CBASP) study; COMPULS/TS EUROTRAIN, European-Wide Investigation and Training Network on the Etiology and Pathophysiology of Gilles de la Tourette Syndrome; GSP, The Brain Genomics Superstruct Project; HCP Aging, Adult Brain, Development, Human Connectome Project; Healthy Brain Network/CMI, Healthy Brain Network/Child Mind Institute; ICBM, International Consortium for Brain Mapping; IMAGEN, The IMAGEN Consortium; IMH, Institute of Mental Health, Singapore; Indiana, Indiana University School of Medicine; IRCCS-FSL, Istituto di Ricovero e Cura a Carattere Scientifico - Fondazione Santa Lucia; Leiden, Leiden University; MAS, Memory and Ageing Study; MCIC, MIND Clinical Imaging Consortium formed by the Mental Illness and Neuroscience Discovery (MIND) Institute, now the Mind Research Network; Melbourne, University of Melbourne; Muenster, Muenster Neuroimaging Cohort; Neuroventure, the imaging part of the Co-Venture Trial funded by the Canadian Institute of Health Research (CIHR); NUIG, National University of Ireland, Galway; NUIG, National University of Ireland, Galway; NYU_TRT, New York University – Test-Retest Reliability; Olin, Olin Neuropsychiatric Research Center; PING, Pediatric Imaging, Neurocognition, and Genetics; QTIM, Queensland Twin Imaging Study; Rockland, Nathan Kline Institute-Rockland Sample; Sydney, University of Sydney; UNIBA, University of Bari Aldo Moro; UPENN, University of Pennsylvania; UTH, University of Texas Health | | | | | | | |
