## supplementary table 2 for "Brain-Age Prediction: Systematic Evaluation of Site Effects, and Sample Age Range and Size"

| **Supplementary Table S2.** **Details of the replication and consistency samples** | | | | | | | |
| --- | --- | --- | --- | --- | --- | --- | --- |
| **Dataset Abbreviation** | **Location** | **Scanner Strength (Tesla)** | **N** | **Age (years) Mean (SD)** | **Age (years) Range** | **Male/Female** | **Further Information and Access Requests** |
| **Replication Sample** | | | | | | | |
| CHCP | China | 3T | 361 | 34.14 (18.22) | 18-79 | 166/195 | http://www.chinese-hcp.cn/ |
| Lexical | USA | 3T | 91 | 11.35 (2.08) | 9-16 | 52/39 | https://openneuro.org/datasets/ds002236/versions/1.1.1 |
| MICA | Canada | 3T | 83 | 31.07 (8.15) | 19-60 | 45/38 | https://osf.io/j532r/ |
| NIMH | USA | 3T | 155 | 34.05 (12.75) | 18-72 | 53/102 | https://openneuro.org/datasets/ds004215/versions/1.0.2 |
| QTAB | Australia | 3T | 396 | 10.84 (1.34) | 8-14 | 205/191 | https://openneuro.org/datasets/ds004146/versions/1.0.4 |
| SALD | China | 3T | 494 | 45.18 (17.44) | 19-80 | 187/307 | http://fcon_1000.projects.nitrc.org/indi/retro/sald.html |
| SLIM | China | 3T | 522 | 20.07 (1.29) | 17-27 | 231/291 | http://fcon_1000.projects.nitrc.org/indi/retro/southwestuni_qiu_index.html |
| **Total** | **-** | **-** | **2102** | **27.74 (17.48)** | **8-80** | **939/1163** | - |
| **Longitudinal Consistency Sample** | | | | | | | |
| QTAB-scan 1 | Australia | 3T | 259 | 10.86 (1.34) | 9-14 | 130/129 | https://openneuro.org/datasets/ds004146/versions/1.0.4 |
| QTAB-scan 2 | Australia | 3T | 259 | 12.53 (1.53) | 10-16 | 130/129 | https://openneuro.org/datasets/ds004146/versions/1.0.4 |
| SLIM-scan 1 | China | 3T | 118 | 19.69 (0.94) | 17-22 | 59/59 | http://fcon_1000.projects.nitrc.org/indi/retro/southwestuni_qiu_index.html |
| SLIM-scan 2 | China | 3T | 118 | 22.07 (0.88) | 19-25 | 59/59 | http://fcon_1000.projects.nitrc.org/indi/retro/southwestuni_qiu_index.html |
| Total | - | - | 377 | 14.57 (4.55) | 9-25 | 189/188 | - |
| CHCP: Chinese Human Connectome Project; Lexical: The Cross-Sectional Multidomain Lexical Processing Dataset; MICA: Microstructure-Informed Connectomics Dataset; NIMH: The National Institute of Mental Health (NIMH) Intramural Healthy Volunteer Dataset; QTAB: The Queensland Twin Adolescent Brain Project; SALD: Southwest University Adult Lifespan Dataset; SLIM: Southwest University Longitudinal Imaging Multimodal Brain Data. | | | | | | | |
