## supplementary table 3 for "Brain-Age Prediction: Systematic Evaluation of Site Effects, and Sample Age Range and Size"

| **Table S3. Females Only. Mean and standard deviation (SD) of the mean absolute error (MAE) and of the correlation coefficient between the chronological age and brain-age in the discovery sample (CORR); Adjusted MAE and Adjusted CORR were corrected for age-bias.** | | | | | | | |
| --- | --- | --- | --- | --- | --- | --- | --- |
| **Age range (years)** | **Mean Chronological Age (SD)** | **MAE (SD)** | **CORR (SD)** | **Adjusted MAE (SD)** | **Adjusted CORR (SD)** | **MAE change with bias correction** | **CORR change with bias correction** |
| 10-year intervals | | | | | |  |  |
| **5≤age<10** | 9.26 (0.63) | 0.37 (0.02) | 0.49 (0.09) | 0.20 (0.02) | 0.90 (0.03) | -45.95% | 83.67% |
| **10≤age<20** | 14.92 (3.42) | 1.12 (0.05) | 0.90 (0.01) | 1.04 (0.03) | 0.93 (0.01) | -7.14% | 3.33% |
| **20≤age<30** | 24.01 (2.71) | 1.80 (0.05) | 0.54 (0.05) | 1.09 (0.06) | 0.89 (0.02) | -39.44% | 64.81% |
| **30≤age<40** | 34.05 (2.61) | 1.61 (0.16) | 0.29 (0.15) | 0.75 (0.13) | 0.90 (0.05) | -53.42% | 210.34% |
| **40≤age<50** | 47.12 (2.89) | 1.71 (0.15) | 0.60 (0.09) | 1.08 (0.12) | 0.90 (0.04) | -36.84% | 50.00% |
| **50≤age<60** | 55.67 (2.82) | 2.33 (0.07) | 0.24 (0.04) | 0.65 (0.03) | 0.96 (0.01) | -72.10% | 300.00% |
| **60≤age<70** | 65.32 (2.84) | 2.28 (0.09) | 0.32 (0.04) | 0.82 (0.06) | 0.94 (0.01) | -64.04% | 193.75% |
| **70≤age<80** | 74.01 (2.59) | 2.00 (0.16) | 0.32 (0.08) | 0.67 (0.04) | 0.95 (0.01) | -66.50% | 196.88% |
| **80≤age≤90** | 83.55 (2.55) | 1.98 (0.30) | 0.36 (0.27) | 0.76 (0.17) | 0.95 (0.04) | -61.62% | 163.89% |
| 20-year intervals | | | | | |  |  |
| **5≤age<20** | 13.24 (3.87) | 1.17 (0.03) | 0.91 (0.01) | 1.1 (0.04) | 0.94 (0.01) | -5.98% | 3.3% |
| **20≤age<40** | 25.77 (4.67) | 2.62 (0.08) | 0.63 (0.03) | 1.89 (0.09) | 0.89 (0.01) | -27.86% | 41.27% |
| **40≤age<60** | 54.15 (4.33) | 2.96 (0.12) | 0.43 (0.04) | 1.34 (0.07) | 0.92 (0.01) | -54.73% | 113.95% |
| **60≤age≤80** | 67.88 (4.84) | 3.13 (0.08) | 0.58 (0.03) | 1.86 (0.07) | 0.9 (0.01) | -40.58% | 55.17% |
| 30-year intervals | | | | | |  |  |
| **5≤age<30** | 17.05 (6.23) | 2.01 (0.08) | 0.9 (0.01) | 1.89 (0.07) | 0.93 (0.01) | -5.97% | 3.33% |
| **30≤age<60** | 51.57 (7.90) | 3.38 (0.09) | 0.81 (0.01) | 2.77 (0.09) | 0.90 (0.01) | -18.05% | 11.11% |
| **60≤age≤90** | 68.50 (5.66) | 3.26 (0.05) | 0.67 (0.03) | 2.17 (0.09) | 0.89 (0.01) | -33.44% | 32.84% |
| 40-year intervals | | | | | |  |  |
| **5≤age<40** | 18.24 (7.44) | 2.38 (0.08) | 0.89 (0.01) | 2.26 (0.05) | 0.93 (0.0) | -5.04% | 4.49% |
| **40≤age≤80** | 61.83 (8.24) | 4.65 (0.1) | 0.68 (0.01) | 3.35 (0.12) | 0.89 (0.01) | -27.96% | 30.88% |
| full age range | | | | | |  |  |
| **5≤age≤90** | 41.34 (23.48) | 5.18 (0.11) | 0.95 (0.00) | 5.09 (0.11) | 0.96 (0.00) | -1.74% | 1.05% |
