## supplementary table 4 for "Brain-Age Prediction: Systematic Evaluation of Site Effects, and Sample Age Range and Size"

| **Table S4. Males Only. Mean and standard deviation (SD) of the mean absolute error (MAE) and of the correlation coefficient (CORR) between the chronological age and brain-age in the discovery sample; Adjusted MAE and Adjusted CORR were corrected for age-bias.** | | | | | | | |
| --- | --- | --- | --- | --- | --- | --- | --- |
| **Age range (years)** | **Mean**  **Chronological**  **Age (SD)** | **MAE (SD)** | **CORR (SD)** | **Adjusted MAE (SD)** | **Adjusted CORR (SD)** | **MAE change with bias correction** | **CORR change with bias correction** |
| 10-year intervals | | | | | | | |
| **5≤age<10** | 9.20 (0.75) | 0.43 (0.04) | 0.58 (0.06) | 0.27 (0.02) | 0.9 (0.02) | -37.21% | 55.17% |
| **10≤age<20** | 14.40 (3.26) | 1.09 (0.05) | 0.88 (0.02) | 1.01 (0.05) | 0.92 (0.01) | -7.34% | 4.55% |
| **20≤age<30** | 24.24 (2.77) | 1.86 (0.17) | 0.5 (0.07) | 1.1 (0.07) | 0.89 (0.02) | -40.86% | 78% |
| **30≤age<40** | 34.18 (2.79) | 1.85 (0.33) | 0.43 (0.16) | 0.96 (0.14) | 0.91 (0.04) | -48.11% | 111.63% |
| **40≤age<50** | 47.06 (2.86) | 1.65 (0.16) | 0.59 (0.08) | 1.03 (0.11) | 0.91 (0.03) | -37.58% | 54.24% |
| **50≤age<60** | 55.54 (2.83) | 2.33 (0.08) | 0.27 (0.05) | 0.72 (0.04) | 0.95 (0.0) | -69.1% | 251.85% |
| **60≤age<70** | 65.40 (2.91) | 2.35 (0.1) | 0.3 (0.06) | 0.84 (0.05) | 0.94 (0.01) | -64.26% | 213.33% |
| **70≤age<80** | 74.10 (2.64) | 2.0 (0.16) | 0.32 (0.08) | 0.67 (0.04) | 0.95 (0.01) | -66.5% | 196.88% |
| **80≤age≤90** | 83.44 (2.61) | 2.03 (0.58) | 0.32 (0.21) | 0.33 (0.14) | 0.98 (0.02) | -83.74% | 206.25% |
| 20-year intervals | | | | | | | |
| **5≤age<20** | 12.95 (3.64) | 1.17 (0.05) | 0.89 (0.01) | 1.09 (0.03) | 0.93 (0.0) | -6.84% | 4.49% |
| **20≤age<40** | 26.29 (4.88) | 3.0 (0.18) | 0.58 (0.05) | 1.9 (0.17) | 0.9 (0.02) | -36.67% | 55.17% |
| **40≤age<60** | 53.65 (4.53) | 2.99 (0.11) | 0.49 (0.05) | 1.54 (0.12) | 0.9 (0.02) | -48.49% | 83.67% |
| **60≤age≤80** | 68.32 (4.99) | 3.31 (0.15) | 0.56 (0.02) | 1.89 (0.08) | 0.9 (0.01) | -42.9% | 60.71% |
| 30-year intervals | | | | | | | |
| **5≤age<30** | 16.47 (6.23) | 1.97 (0.06) | 0.9 (0.0) | 1.87 (0.03) | 0.93 (0.0) | -5.08% | 3.33% |
| **30≤age<60** | 50.76 (8.16) | 3.48 (0.15) | 0.8 (0.01) | 2.83 (0.1) | 0.9 (0.01) | -18.68% | 12.5% |
| **60≤age≤90** | 68.93 (5.75) | 3.47 (0.11) | 0.63 (0.04) | 2.21 (0.12) | 0.89 (0.02) | -36.31% | 41.27% |
| 40-year intervals | | | | | | | |
| **5≤age<40** | 17.79 (7.63) | 2.45 (0.07) | 0.89 (0.01) | 2.32 (0.04) | 0.93 (0.0) | -5.31% | 4.49% |
| **40≤age≤80** | 61.95 (8.71) | 4.83 (0.15) | 0.7 (0.02) | 3.56 (0.1) | 0.88 (0.01) | -26.29% | 25.71% |
| full age range | | | | | | | |
| **5≤age≤90** | 40.23 (23.88) | 5.21 (0.08) | 0.96 (0.00) | 5.12 (0.07) | 0.96 (0.00) | -1.73% | 0.00% |
