## supplementary table 5 for "Brain-Age Prediction: Systematic Evaluation of Site Effects, and Sample Age Range and Size"

| **Table S5. Females Only. Mean and standard deviation (SD) of the mean absolute error (MAE) and of the correlation coefficient (CORR) between the chronological age and brain-age in the replication sample; Adjusted MAE and Adjusted CORR were corrected for age-bias.** | | | | | | | |
| --- | --- | --- | --- | --- | --- | --- | --- |
| **Age range (years)** | **Mean**  **Chronological Age (SD)** | **MAE** | **CORR** | **Adjusted MAE** | **Adjusted CORR** | **MAE change with bias correction** | **CORR change with bias correction** |
| 10-year intervals | | | | | | | |
| **5≤age<10** | 9.42 (0.53) | 0.55 | 0.15 | 0.29 | 0.85 | -47.27% | 466.67% |
| **10≤age<20** | 16.96 (3.59) | 2.64 | 0.51 | 2.06 | 0.72 | -21.97% | 41.18% |
| **20≤age<30** | 23.59 (2.41) | 2.25 | 0.03 | 1.19 | 0.82 | -47.11% | 2633.33% |
| **30≤age<40** | 35.00 (2.56) | 2.14 | 0.25 | 1.47 | 0.91 | -31.31% | 264.00% |
| **40≤age<50** | 46.09 (2.76) | 2.67 | 0.26 | 1.18 | 0.90 | -55.81% | 246.15% |
| **50≤age<60** | 56.45 (2.82) | 2.43 | 0.08 | 0.61 | 0.97 | -74.90% | 1112.50% |
| **60≤age<70** | 64.05 (2.65) | 2.59 | 0.34 | 0.79 | 0.95 | -69.50% | 179.41% |
| **70≤age≤80** | 73.89 (1.83) | 1.54 | 0.14 | 0.65 | 0.91 | -57.79% | 550.00% |
| 20-year intervals | | | | | | | |
| **5≤age<20** | 15.70 (4.32) | 2.74 | 0.66 | 2.26 | 0.78 | -17.52% | 18.18% |
| **20≤age<40** | 25.57 (4.96) | 3.80 | 0.34 | 2.09 | 0.87 | -45.00% | 155.88% |
| **40≤age<60** | 52.45 (5.77) | 4.69 | 0.31 | 1.42 | 0.95 | -69.72% | 206.45% |
| **60≤age≤80** | 66.71 (5.01) | 11.50 | 0.40 | 2.17 | 0.95 | -81.13% | 137.50% |
| 30-year intervals | | | | | | | |
| **5≤age<30** | 18.58 (5.34) | 3.25 | 0.73 | 3.01 | 0.82 | -7.38% | 12.33% |
| **30≤age<60** | 47.91 (9.24) | 7.07 | 0.43 | 4.79 | 0.76 | -32.25% | 76.74% |
| **60≤age≤90** | 66.71 (5.03) | 4.87 | 0.35 | 2.89 | 0.83 | -40.66% | 137.14% |
| 40-year intervals | | | | | | | |
| **5≤age<40** | 19.75 (6.69) | 3.91 | 0.71 | 3.53 | 0.83 | -9.72% | 16.90% |
| **40≤age≤80** | 57.56 (8.78) | 6.95 | 0.63 | 4.45 | 0.86 | -35.97% | 36.51% |
| full age range | | | | | | | |
| **5≤age≤90** | 28.82 (17.70) | 8.33 | 0.86 | 7.30 | 0.88 | -12.36% | 2.33% |
