## supplementary table 6 for "Brain-Age Prediction: Systematic Evaluation of Site Effects, and Sample Age Range and Size"

| **Table S6. Males Only. Mean and standard deviation (SD) of the mean absolute error (MAE) and of the correlation coefficient (CORR) between the chronological age and brain-age in the replication sample; Adjusted MAE and Adjusted CORR were corrected for age-bias.** | | | | | | | |
| --- | --- | --- | --- | --- | --- | --- | --- |
| **Age range (years)** | **Mean**  **Chronological Age (SD)** | **MAE** | **CORR** | **Adjusted MAE** | **Adjusted CORR** | **MAE change with bias correction** | **CORR change with bias correction** |
| 10-year intervals | | | | | | | |
| **5≤age<10** | 9.36 (0.52) | 0.53 | 0.18 | 0.35 | 0.71 | -33.96% | 294.44% |
| **10≤age<20** | 15.95 (3.92) | 3.02 | 0.50 | 2.19 | 0.77 | -27.48% | 54.00% |
| **20≤age<30** | 23.68 (2.61) | 2.39 | 0.11 | 1.26 | 0.84 | -47.28% | 663.64% |
| **30≤age<40** | 34.71 (2.59) | 2.21 | 0.35 | 1.36 | 0.92 | -38.46% | 162.86% |
| **40≤age<50** | 45.40 (3.04) | 3.15 | 0.11 | 1.59 | 0.82 | -49.52% | 645.45% |
| **50≤age<60** | 56.73 (2.72) | 2.54 | 0.18 | 0.79 | 0.94 | -68.90% | 422.22% |
| **60≤age<70** | 64.37 (2.72) | 2.70 | 0.32 | 0.89 | 0.95 | -67.04% | 196.88% |
| **70≤age≤80** | 74.85 (2.41) | 2.04 | 0.10 | 0.65 | 0.95 | -68.14% | 850.00% |
| 20-year intervals | | | | | | | |
| **5≤age<20** | 14.45 (4.42) | 3.04 | 0.63 | 2.31 | 0.80 | -24.01% | 26.98% |
| **20≤age<40** | 25.60 (4.95) | 3.93 | 0.40 | 2.20 | 0.87 | -44.02% | 117.50% |
| **40≤age<60** | 52.90 (6.06) | 4.82 | 0.32 | 1.94 | 0.92 | -59.75% | 187.50% |
| **60≤age≤80** | 67.09 (5.30) | 12.80 | 0.30 | 3.61 | 0.91 | -71.80% | 203.33% |
| 30-year intervals | | | | | | | |
| **5≤age<30** | 18.01 (5.91) | 3.23 | 0.76 | 2.91 | 0.85 | -9.91% | 11.84% |
| **30≤age<60** | 45.66 (10.24) | 8.30 | 0.49 | 5.38 | 0.80 | -35.18% | 63.27% |
| **60≤age≤90** | 67.09 (5.34) | 4.57 | 0.46 | 2.59 | 0.86 | -43.33% | 86.96% |
| 40-year intervals | | | | | | | |
| **5≤age<40** | 19.28 (7.24) | 4.02 | 0.74 | 3.60 | 0.84 | -10.45% | 13.51% |
| **40≤age≤80** | 59.49 (9.10) | 6.24 | 0.63 | 4.09 | 0.87 | -34.46% | 38.10% |
| full age range | | | | | | | |
| **5≤age≤90** | 26.39 (17.12) | 8.71 | 0.85 | 7.52 | 0.87 | -13.66% | 2.35% |
