## supplementary table 7 for "Brain-Age Prediction: Systematic Evaluation of Site Effects, and Sample Age Range and Size"

| **Table S7. Pre-trained model performances in the independent longitudinal-consistency sample with different age ranges of female participants. MAE: mean absolute error; CORR: correlation coefficient between the chronological age and predicted brain age.** | | | | | | | | | | | | | | |
| --- | --- | --- | --- | --- | --- | --- | --- | --- | --- | --- | --- | --- | --- | --- |
|  | **Scan 1** | | | | | | | **Scan 2** | | | | | | |
| **Age range (years)** | **Mean age (SD)** | **MAE** | **CORR** | **Adjusted MAE** | **Adjusted CORR** | **MAE change with bias correction** | **CORR change with bias correction** | **Mean age (SD)** | **MAE** | **CORR** | **Adjusted MAE** | **Adjusted CORR** | **MAE change with bias correction** | **CORR change with bias correction** |
| 10-year intervals | | | | | | | | | | | | | | |
| **5≤age<10** | 8.94 (0.25) | 0.52 | 0.37 | 0.28 | 0.90 | -46.15% | 143.24% | 10 (0) | 1.10 | n/a | 0.58 | n/a | -47.27% | n/a |
| **10≤age<20** | 11.94 (1.06) | 2.84 | 0.59 | 2.29 | 0.78 | -19.37% | 32.2% | 13.78 (1.14) | 1.83 | 0.65 | 1.57 | 0.73 | -14.21% | 12.31% |
| **20≤age<30** | 21.20 (0.42) | 2.48 | 0.10 | 0.96 | 0.45 | -61.29% | 350.00% | 23.53 (0.51) | 1.27 | 0.17 | 0.52 | 0.78 | -59.06% | 358.82% |
| 20-year intervals | | | | | | | | | | | | | | |
| **5≤age<20** | 11.00 (1.51) | 2.72 | 0.68 | 2.37 | 0.79 | -12.87% | 16.18% | 12.72 (1.71) | 1.67 | 0.71 | 1.71 | 0.77 | 2.4% | 8.45% |
| **20≤age<40** | 21.20 (0.42) | 4.66 | 0.09 | 2.20 | 0.24 | -52.79% | 166.67% | 23.53 (0.51) | 3.46 | 0.14 | 1.76 | 0.41 | -49.13% | 192.86% |
| 30-year intervals | | | | | | | | | | | | | | |
| **5≤age<30** | 13.66 (4.22) | 3.83 | 0.68 | 3.25 | 0.78 | -15.14% | 14.71% | 15.59 (4.56) | 2.71 | 0.75 | 2.57 | 0.82 | -5.17% | 9.33% |
| 40-year intervals | | | | | | | | | | | | | | |
| **5≤age<40** | 13.66 (4.22) | 4.13 | 0.70 | 3.39 | 0.79 | -17.92% | 12.86% | 15.59 (4.56) | 2.97 | 0.77 | 2.84 | 0.84 | -4.38% | 9.09% |
| full age range | | | | | | | | | | | | | | |
| **5≤age≤90** | 13.66 (4.22) | 12.70 | 0.59 | 10.60 | 0.61 | -16.54% | 3.39% | 15.59 (4.56) | 13.00 | 0.72 | 11.10 | 0.74 | -14.62% | 2.78% |
