## supplementary table 8 for "Brain-Age Prediction: Systematic Evaluation of Site Effects, and Sample Age Range and Size"

| **Table S8. Pre-trained model performances in the independent longitudinal-consistency sample with different age ranges of male participants. MAE: mean absolute error; CORR: correlation coefficient between the chronological age and predicted brain age.** | | | | | | | | | | | | | | |
| --- | --- | --- | --- | --- | --- | --- | --- | --- | --- | --- | --- | --- | --- | --- |
|  | **Scan 1** | | | | | | | **Scan 2** | | | | | | |
| **Age range (years)** | **Mean age (SD)** | **MAE** | **CORR** | **Adjusted MAE** | **Adjusted CORR** | **MAE change with bias correction** | **CORR change with bias correction** | **Mean age (SD)** | **MAE** | **CORR** | **Adjusted MAE** | **Adjusted CORR** | **MAE change with bias correction** | **CORR change with bias correction** |
| 10-year intervals | | | | | | | | | | | | | | |
| **5≤age<10** | 8.94 (0.25) | 0.52 | 0.26 | 0.30 | 0.79 | -42.31% | 203.85% | 10.00 (0) | 1.22 | n/a | 0.71 | n/a | -41.8% | n/a |
| **10≤age<20** | 11.61 (0.75) | 3.19 | 0.47 | 2.35 | 0.77 | -26.33% | 63.83% | 13.29 (0.96) | 1.94 | 0.58 | 1.66 | 0.68 | -14.43% | 17.24% |
| **20≤age<30** | 21.17 (0.39) | 2.52 | 0.42 | 1.12 | 0.23 | -55.56% | -45.24% | 23.33 (0.35) | 1.11 | 0.18 | 1.05 | 0.59 | -5.41% | 227.78% |
| 20-year intervals | | | | | | | | | | | | | | |
| **5≤age<20** | 10.78 (1.26) | 3.11 | 0.55 | 2.45 | 0.77 | -21.22% | 40.00% | 12.40 (1.44) | 1.56 | 0.65 | 1.55 | 0.72 | -0.64% | 10.77% |
| **20≤age<40** | 21.17 (0.39) | 5.46 | 0.28 | 2.89 | 0.18 | -47.07% | -35.71% | 23.33 (0.35) | 3.33 | 0.15 | 1.78 | 0.38 | -46.55% | 153.33% |
| 30-year intervals | | | | | | | | | | | | | | |
| **5≤age<30** | 13.60 (4.35) | 3.44 | 0.60 | 2.82 | 0.74 | -18.02% | 23.33% | 15.45 (4.71) | 2.60 | 0.74 | 2.28 | 0.82 | -12.31% | 10.81% |
| 40-year intervals | | | | | | | | | | | | | | |
| **5≤age<40** | 13.60 (4.35) | 3.93 | 0.61 | 3.19 | 0.74 | -18.83% | 21.31% | 15.45 (4.71) | 2.91 | 0.72 | 2.71 | 0.81 | -6.87% | 12.50% |
| full age range | | | | | | | | | | | | | | |
| **5≤age≤90** | 13.60 (4.35) | 12.00 | 0.56 | 9.94 | 0.59 | -17.17% | 5.36% | 15.45 (4.71) | 11.50 | 0.74 | 9.83 | 0.75 | -14.52% | 1.35% |
